# Exploring spatio-temporal neural dynamics of the human visual cortex

**DOI:** 10.1101/422576

**Authors:** Ying Yang, Michael J. Tarr, Robert E. Kass, Elissa M. Aminoff

## Abstract

The human visual cortex is organized in a hierarchical manner. Although a significant body of evidence has been accumulated in support of this hypothesis, specific details regarding the spatial and temporal information flow remain open. Here we present detailed spatio-temporal correlation profiles of neural activity with low-level and high-level features derived from a “deep” (8-layer) neural network pre-trained for object recognition. These correlation profiles indicate an early-to-late shift from low-level features to high-level features and from low-level regions to higher-level regions along the visual hierarchy, consistent with feedforward information flow. To refine our understanding of information flow, we computed three sets of features from the low-and high-level features provided by the neural network: object-category-relevant low-level features (the common components between low-level and high-level features), low-level features roughly orthogonal to high-level features (the residual Layer 1 features), and unique high-level features that were roughly orthogonal to low-level features (the residual Layer 7 features). Contrasting the correlation effects of the common components and the residual Layer 1 features, we observed that the early visual cortex exhibits a similar amount of correlation with the two feature sets early in time (60 to 120 ms), but in a later time window, the early visual cortex exhibits a higher and longer correlation effect with the common components/low-level task-relevant features as compared to the low-level residual features—an effect unlikely to arise from purely feedforward information flow. Overall, our results indicate that non-feedforward processes, for example, top-down influences from mental representations of categories, may facilitate differentiation between these two types of low-level features within the early visual cortex.

## 1 Introduction

Humans can effortlessly recognize objects and understand scenes. What computational processes in the visual cortex result in such proficiency? Decades of research has mapped visual cortex into multiple functional brain areas, which span those defined through a topological mapping (i.e., “retinotopy”) of the visual field (Engel et al., 1997) and those defined through preferred stimulus category (e.g., scene-selective parahippocampal place area, Epstein et al. (1999) and object-selective “lateral occipital complex” Grill-Spector et al. (1999)). More generally, the visual cortex is hypothesized to process visual inputs in a hierarchical manner (DiCarlo and Cox, 2007). At a low level, neurons in primary visual cortex (V1) have small receptive fields and extract low-level information such as local edge orientations (Hubel and Wiesel, 1968). Next in the hierarchy, neurons in V2 are sensitive to somewhat more complex edge angles and junctions (Ito and Komatsu, 2004), as well as textures (Freeman et al., 2013). This pattern continues, with regions of the visual cortex progressively processing more and more complex visual representations – ultimately resulting in high-level vision. For example, neurons in the inferior temporal cortex show selectivity to complex shapes, invariance to scales and lighting, and encode semantic information associated with visual inputs (Tanaka, 1996). To be clear, there is a very extensive body of experimental evidence supporting this hierarchical organization, ranging from electrophysiology studies in primates to neuroimaging studies in humans (e.g., Yamins et al. (2014); Cichy et al. (2016b); Aminoff et al.(2015); Clarke et al. (2014), etc).

Within the framework of a progressive visual hierarchy, as one moves from the posterior regions (e.g., V1) to the anterior regions along the visual pathways (e.g., category-selective regions), neurons at each level of the hierarchy receive inputs from previous levels and extract increasingly higher-level information. At the same time, we should ask, “Does information necessarily flow only in a bottom-up, feedforward direction?” Apparently not – there is also strong evidence for *feedback* anatomical connections between visual areas, that is connections that pass information in an inverse direction with respect to the hierarchy. Similarly, there is evidence for anatomical connections skipping across levels (e.g., V4 to V1), as well as from the frontal and parietal lobes to visual areas (Felleman and Van Essen, 1991). These various connections indicate that non-feedforward dynamics are likely to exist and, moreover, that this sort of information flow may function as top-down feedback, thereby facilitating rapid visual processing and recognition (e.g., as suggested by Bar et al. (2006)).

Although there is some functional evidence for top-down information flow in visual cortex, much of the support for feedback is neuroanatomical. As such, further study of the spatio-temporal dynamics of visual processing is critical. For example, to address the directionality of information flow when visual inputs are processed, it is important to know not only where information is processed, but when it is processed by one brain region relative to other brain regions. That is, one should be able to explicate the level of information extracted at a given time point from a specific brain region and compare this to information extracted from other regions at other time points. Such data would inform us regarding whether information flows in the direction of the apparent feedback (i.e., not only in a feedforward direction when processing visual inputs). Moreover, if evidence for top-down processing is found, when and in what brain areas does such feedback occur? Details of information flow dynamics are crucial for building mathematical models of the brain and understanding how visual recognition is achieved.

To address such questions we need to better understand neural dynamics in the context of the neural activity associated with the perception of a rich set of visual stimuli (e.g., naturalistic, everyday visual scenes). Moreover, the ideal measurement tool will provide both good temporal and good spatial resolution. With respect to the former, magnetoencepholography (MEG) offers millisecond-level temporal resolution, enabling the measurement of rapid neural processing in object and scene recognition. With respect to the latter, MEG also offers reasonable source localization techniques (e.g., Hamalainen et al. (1993); Dale et al. (2000);Yang et al. (2014)), enabling the localization of neural responses at an intermediate spatial resolution. Together, the two characteristics of MEG supports a relatively detailed spatio-temporal profile of neural activity. As such, we have adopted MEG as a promising neuroimaging method for non-invasively studying neural activity in response to a visual processing paradigm aimed at addressing the question—*what is the form of information extracted at different temporal stages and different spatial locations during scene and object perception*?

To address this question, it is critical to use MEG strengths (e.g., spatial and temporal resolution) to explore the level of information representation (e.g., edges vs. full shapes vs. meaningful objects) in the spatio-temporal dynamics of neural processing. To accomplish this, one can regress measured spatio-temporal neural activity during a visual task against different candidate feature sets that characterize different levels of information (Nestor et al., 2008). However, defining the candidate features is non-trivial. Building on a tradition of applying computer vision models to the understanding of biological vision (Leeds et al., 2013), many different groups have recently used a type of artificial-neural-network model, known as convolutional neural networks (”CNNs”) to study human visual processing (Yamins and DiCarlo, 2016). What makes CNNs particularly attractive as “proxy models” for studying biological vision is their extremely high performance in categorizing both naturalistic objects and scenes, as well as their demonstrated ability to learn task-relevant, diagnostic features across very large image sets (Krizhevsky et al., 2012). Interestingly, CNNs (and related “deep” networks) were inspired by feedforward processing as embodied in the hypothesized hierarchical structure of the primate visual cortex (LeCun et al., 2015). In particularly, CNNs typically incorporate multiple layers with units of increasing receptive field sizes as one moves up the hierarchy; the first layer takes raw image inputs and the last layer outputs object or scene category labels. The connections between layers are optimized by minimizing labeling error on an extremely large amount of image data (1,000,000’s of examples). After such optimization or training, the layered structure of the network inherently provides an operational definition of low-level to high-level representation of task-relevant visual information (Zeiler and Fergus, 2014). Moreover, as alluded to earlier, features extracted from CNNs have been found to share significant representational similarity with the neural representations of objects and scenes at both low levels, as exemplified by early visual regions, and high levels, as exemplified by category-selective brain regions (Yamins et al., 2014; Khaligh-Razavi and Kriegeskorte, 2014; Cichy et al., 2016a,c).

In this context, we used CNNs to explicate the level of visual representation within a spatio-temporal context. To the degree that we can use this approach to associate low-and high-level visual features with specific processing time points, we can develop a more detailed account of feedforward versus feedback communication within visual cortex. More specifically, we regressed the neural responses — as measured by MEG — at different cortical locations and time points against low-and high-level features from a high-performing CNN. Critically, this regression analysis explores how well neural responses are explained by particular feature sets, yielding a spatial-temporal profile of statistical dependence between neural activity and different levels of CNN (or some other model) features. In the ideal case, using an unlimited number of observations of neural responses and an unlimited number of naturalistic stimuli, we could adopt a high-capacity regression model to capture statistical dependence. However, the number of observations we can actually acquire is limited by our experimental methods; therefore, we use linear regression to restrict model complexity and avoid overfitting. In this context, the spatio-temporal profile of linear dependence, which we refer to as the “correlation profile,” is the key result we present in this paper. More specifically, the spatio-temporal correlation profile sheds light on information flow within the visual cortex.

In this same spirit, several recent MEG studies have likewise related neural activity to CNN or other computer vision derived features. However, in contrast to our present study, these studies focused on recognizing isolated objects, that is, restricted to individual objects on blank backgrounds (Clarke et al., 2014; Cichy et al., 2016c), or on specific properties of scenes, that is, restricting the input to a small set of scene stimuli (Cichy et al., 2016a). For the most part, these studies also focused only on temporal patterns in MEG, and not on spatial patterns (although see Clarke et al. (2014)). To address what we view as limitations, we recorded neural responses to a relatively large number of *everyday, naturalistic scenes* and applied *source-space regression analysis*, aiming to obtain detailed correlation profiles between spatio-temporal neural activity and low-and high-level CNN features.

Beyond these extensions of prior work, our present study is distinguished by our implementation of a unique decomposition of low-level and high-level CNN features. Because the low-and high-level features from CNNs are defined empirically by large-scale learning as instantiated within the neural network, high-level features are non-linear transforms of low-level features. As such, features at different levels may share common linear components. The existence of these common components renders it difficult to interpret correlations between neural data and model features at different levels (as has been done previously); that is, correlations across comparisons may be driven by common components. In response to this challenge, we present a new approach, which extracts the common components in the low-and high-level CNN features and decomposes them into three groups:

1. *Common components* between the low-level and high-level features, or the subspace of low-level features that are relevant to high-level features.
2. *Residuals of low-level features.* These are the low-level features that are roughly orthogonal to high-level features, or the features remaining after regressing out the common components.
3. *Residuals of high-level features.* These are the high-level features that are roughly orthogonal to low-level features, or the features remaining after regressing out the common components.

By comparing the spatio-temporal correlation profiles of these three groups, we are able to observe dynamics for feedforward processing in the visual cortex. Additionally, we see evidence for non-feedforward processing, which we hold may reflect top-down feedback that is deployed as part of naturalistic scene processing (Bar et al., 2006).

## 2 Materials and methods

### 2.1 Participants

Eighteen healthy adults (8 females/10 males, 1 left-handed/17 right-handed, age range 18-30) participated in both the MEG and the fMRI sessions. All participants provided written informed consent and were financially compensated for their participation. All procedures followed the principles in the Declaration of Helsinki and were approved by the Institutional Review Boards of Carnegie Mellon University and the University of Pittsburgh.

### 2.2 Stimuli

The stimulus set used for both MEG and fMRI included color images (photographs) of 181 scene categories (e.g., “airport”, “beach”, “coffee shop”, etc.). There were two exemplar images for each category, resulting in 362 images in total. These images were obtained from the dataset used by the “Never Ending Imaging Learner” (NEIL, Chen et al. (2013), www.neil-kb.com), which were originally scraped from the Internet. The scene images varied in aspect ratio with the longest dimension (either in width or in height) set to 500 pixels. The images were placed in the center of a 600×600 bounding box filled with a gray value of = 135 (out of 255). Figure 1 shows two example stimuli. The images (with the gray bounding boxes) were presented at a visual angle of approximately 10 by 10 degrees.

**Figure 1:**
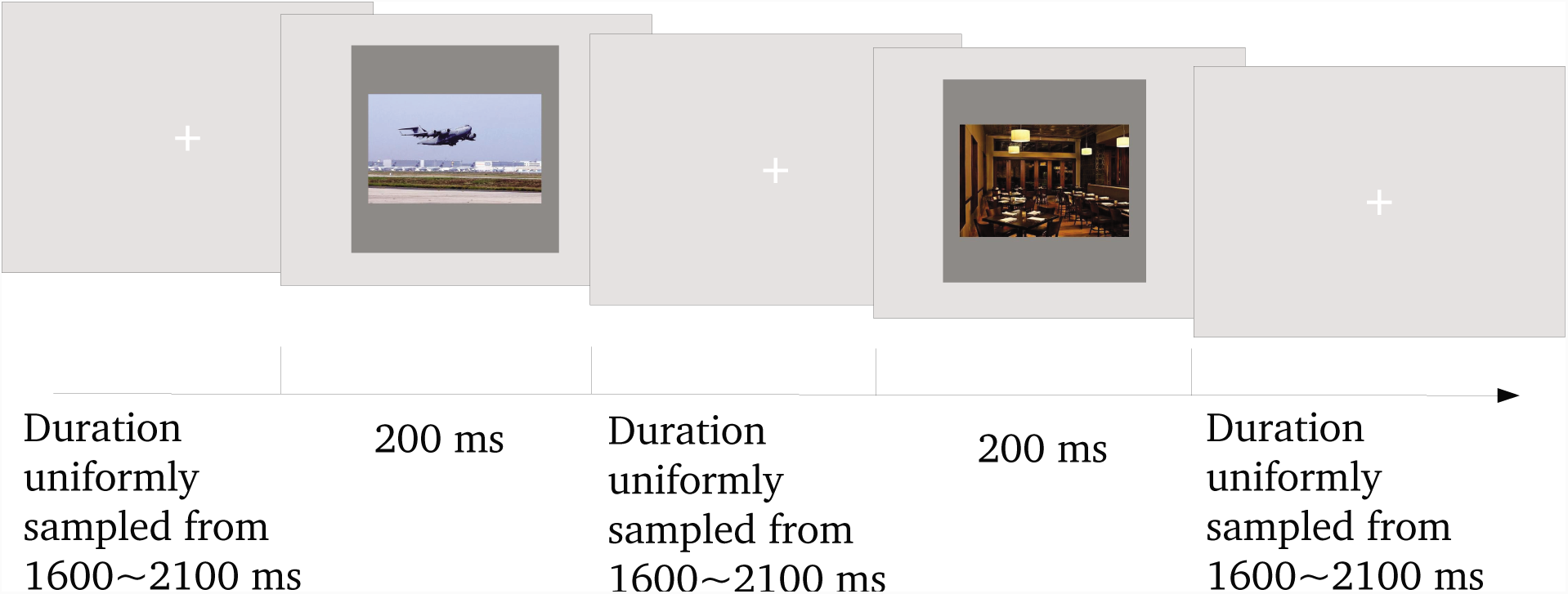
Illustration of the trial structure in the MEG session

In the fMRI session (see below), we also included a functional localizer scanning run in which a separate set of images were used. This localizer was added in order to independently define the object- and scene-selective regions of the visual cortex. The localizer images included 60 color images in each of the three conditions: scenes, objects, and phase-scrambled pictures of the scenes. The scene images included outdoor and indoor scenes, which did not overlap with the 362 scene images discussed above. The particular objects used were weak contextual objects as described in (Bar and Aminoff, 2003). The phase-scrambled pictures served as control stimuli for scenes; they were generated by running a Fourier transform of each scene image, scrambling the phases, and then performing an inverse Fourier transform back into pixel space. The localizer images were presented at a visual angle of 5.5 by 5.5 degrees.

### 2.3 Experimental procedure

The experimental procedures in the MEG and MRI sessions were implemented using `Psychtoolbox 3` (http://psychtoolbox.org/) in `MATLAB`. The trial structure in the MEG sessions is illustrated in Figure 1. Before the stimulus presentation, the participants were asked to fixate their eyes on a white “+” symbol (spanning 80 pixels or 1.3 degrees at the center of the screen. The gray value of the screen was 180 (out of 255). We term this screen “the fixation screen” hereafter. Following the fixation screen, one stimulus image was presented at the center of the screen for 200 ms, which was short enough to reduce artifacts due to saccades during stimulus presentation. The stimulus image was then followed by the fixation screen lasting for a random duration until the onset of the next stimulus. This duration was uniformly sampled from a (1600, 2100) ms interval independently on each trial. Participants performed a “one-back” task — that is, they responded, by pressing a button on a glove pad using the right index finger, when the current stimulus was a repetition of the stimulus from the immediately preceding trial. Participants were given 1500 ms to respond after stimulus onset.

Each MEG session included 6 to 12 runs; each run included 181 images, and every two runs covered all 362 images. In each run, 10% of the images were immediately repeated once, and only the data corresponding to the first presentation of each image were analyzed. In total there were 3 to 6 repetitions of each image for most of the participants in the MEG session. For one participant, due to a problem with acquisition hardware, only 2 repetitions were presented for some images, and 4 repetitions were presented for the remaining images.

Each fMRI session included a functional localizer to independently define scene-and object-selective regions in the brain. Most participants went through two runs of the localizer experiment; however, two participants went through one run due to time limitations. Each run started and ended with a 12-second time window, during which a black fixation cross (“+”) was presented on a gray background. Between the starting and ending fixation windows, there were twelve 14-second to 16-second blocks, each of which presented stimuli in one condition (scenes, weak-contextual objects, or phase-scrambled scenes). There were four blocks per condition and three conditions in total; there was also a 10-second fixation time window between two consecutive blocks. In each block, 14 stimuli were presented in a row, with an 800 ms presentation duration and a 200 ms inter-stimulus interval, with the exception that the first stimulus in each of the block other than the first block was presented for 2800 ms, yielding 16-second blocks Note that the longer, 16-second presentation was due to a timing issue in the customized image presentation program. However, we are doubtful that this variation changed our results in defining the object/scene-selective regions. In particular, we relied on a block design in which the main effect is the difference of neural responses for different blocked conditions. As such, these results are expected to be robust to small variations in the presentation duration of individual stimuli. Among the 14 stimuli, 12 were unique images, and 2 were immediate repetitions of the previous image in order to allow for positive responses in the one-back task — pressing a button on a glove pad with the right index finger when there was an immediate image repetition.

### 2.4 Data acquisition

**MEG.** MEG data was collected using a 306-channel whole-head MEG system (Elekta Neuromag, Helsinki, Finland) at the Brain Mapping Center at the University of Pittsburgh. The MEG system had 102 triplets, each consisting of a magnetometer and two perpendicular gradiometers. The recordings were acquired at 1 kHz, high-pass filtered at 0.1 Hz and low-pass filtered at 330 Hz. Four head position indicator (HPI) coils were placed on the scalp of each participant to record the position of the head in relation to the MEG helmet. Empty room MEG recordings were collected in the same session, and used to estimate the covariance matrix of the sensor noise. Approximately one hundred points describing the shape of the head and the coordinates of the HPI coils on the scalp were collected using a digitization device; these coordinates were later used in aligning the head position in the MEG session with the structural MRI scan.

Electrooculogram (EOG) was monitored by recording electric potentials above and below one eye and lateral to both eyes; electrocardiography (ECG) was recorded by placing additional electrodes under the clavicles. The EOG and ECG recordings captured eye blinks and heartbeats in order to allow these artifacts to be removed from the MEG recordings during data pre-processing.

**MRI.** Magnetic resonance imaging (MRI) data was collected on a 3T Siemens Verio MR scanner at the Scientific Imaging and Brain Research Center at Carnegie Mellon University using a 32-channel head coil. First, for each participant, a high resolution structural MRI scan was acquired (T1-weighted MPRAGE sequence, 1 mm ×1 mm ×1mm, 176 sagittal slices, TR = 2300 ms, TE = 1970 ms, flip angle = 9°, GRAPPA = 2, field of view = 256). Second, functional MRI (fMRI) data was collected for the functional localizer (T2*-weighted echo-planar imaging multiband pulse sequence, 69 slices aligned to the AC/PC, in-plane resolution 2 mm ×2 mm, 2 mm slice thickness, no gap, TR = 2000 ms, TE = 30 ms, flip angle = 79 ° multi-band acceleration factor = 3, field of view 192 mm, phase encoding direction A≫P, ascending acquisition). Third, a fieldmap scan was acquired to correct for distortion effects using the same slice prescription as the echo-planar imaging scans (69 slices aligned to the AC/PC, in-plane resolution 2 mm ×2 mm, 2 mm slice thickness, no gap, TR = 724 ms, TE1 = 5 ms; TE2 = 7.46 ms, flip angle = 70°, field of view 192 mm, phase encoding direction A ≫ P, interleaved acquisition).

### 2.5 Preprocessing of MEG data

Preprocessing of the raw MEG data was accomplished via the following pipeline. All steps in this pipeline were implemented using the MNE-python (Gramfort et al., 2014b) package in Python.

1. *Filtering.* The raw recordings (including MEG empty room recordings) were filtered with a 1-110 Hz bandpass filter, which removed low-frequency drifts and higher frequency noise (such as the oscillations generated by the head position indicator coils during head tracking). Recordings were further filtered by a notch filter centered at 60 Hz intended to remove power line interference.
2. *Removing artifacts due to eye blinks and heartbeats.* Independent component analysis (ICA) was used to decompose the recordings into multiple components. When reconstructing the data from the components, those highly correlated with eye blinks and heartbeats (as recorded by EOG and ECG) were removed.
3. *Obtaining trial-by-trial data.* Trial-by-trial recordings (also referred as “epochs”) were obtained by segmenting the data from −100 ms to 900 ms with respect to the “trigger” onset (the “stimulus onset” as recorded in the acquisition system, defined as 0 ms). For each trial and each channel, the mean across time points in the baseline window (−100 to 0 ms) was subtracted from the recording at each time point. Note that the timing here was recorded by the data acquisition device; the image presentation device showed a measured delay of 40 ms according to a photosensor placed on the screen. As such, to correctly align the data, we shifted all time points backwards by 40 ms. A signal space projection (SSP) was applied to all epochs. The SSP constructed a low-dimensional linear subspace characterizing the empty room noise (via principal component analysis), and removed the projection onto this subspace from the experimental MEG recordings. As such, only those neural signals orthogonal to the principal components of empty room noise remained.
4. *Obtaining averaged neural responses to each image for each participant.* To reduce the computational cost for some of the analyses discussed below, we downsampled the trial-by-trial data to 100 Hz sampling rate. In addition, those trials corresponding to the second presentation in immediate repetitions may have had lower signal strength due to adaptation; therefore they were removed from further analysis. To remove outlier trials that had extreme large variations in each session for each participant, we computed the difference between the maximum and minimum of the recordings for each channel in each trial, and discarded those trials where the difference was larger than 15 standard deviations plus the mean across all trials for at least one channel. Finally, the data in the remaining trials that corresponded to the same image were averaged within each session for each participant.
5. *Regressing out neural data explained by nuisance covariates.* Although our stimulus images were all displayed in the same 60×600 pixel boxes, the widths and heights of the images themselves varied. Such nuisance covariates are irrelevant to the image contents, but could explain a significant amount of variance in the MEG recordings. To remove such spurious effects, we regressed the MEG data against four covariates — image width, image height, area (width×height) and aspect ratio (width */* height) — respectively at each time point for each sensor in each participant. An all-one column was added to the regressors to remove the mean response across all images as well. The residuals were then retained as new sensor data to be analyzed.

### 2.6 Forward modeling

For each participant, based on the T1-weighted structural MRI scan, the outer skin surface of the scalp and the inner and outer surfaces of the skull were computed using the watershed algorithm (Ségonne et al., 2004) implemented in the `Freesurfer` software0020 (Fischl et al., 2002) and the `MNE-C` software (https://martinos.org/mne/dev/install_mne_c.html). Additionally, the cortical surfaces that segmented the gray and white matter were also computed using `Freesurfer`. The source space was defined as about 8000 distributed “dipoles” (or source points) that pointed perpendicularly to the cortical surface of both hemispheres. The average spacing between source points was 4.9 mm, yielding 24 mm^2^ of cortical surface area per source point. Source points that were within 2.5 mm of the inner skull surface were excluded.

The digitized points that described the shape of the head in MEG were used for co-registration with the structural MRI. In the co-registration, we solved for a rigid-body transformation that minimized the sum of squared distances from the digitized points to the scalp surface, using an interface implemented in MNE-C (Gramfort et al., 2014a). Note that the optimization problem was not necessarily convex, therefore no global minimum could be guaranteed. Yet, by manually adjusting initial values, the solution for each participant appeared to be correct in visual inspection.

For each participant, the forward matrix for each run in the MEG session was computed using the boundary element model implemented in `MNE-C`, after transforming the MEG sensor locations into the structural MRI space, based on the alignment in the co-registration step above. The forward matrices across all runs were averaged for each participant.

### 2.7 Source localization and Source-space regression analysis

The dynamic statistical parametric mapping (dSPM) (Dale et al., 2000) source localization method implemented in MNE-python was used to obtain unit-less (i.e., standardized) source current dipole estimates. This method estimates a linear projection that projects the MEG sensor data into the source space. Since there were more source points on the cortical surface maps than the number of sensors, a penalization of the *L*_2_ norm of the linear projection weights was used. The penalization parameter was set to 1.0 in MNE-python. The noise covariance matrix was estimated from sensor recordings within the baseline time windows (−140 to −90 ms) for each participant.

To characterize how much the spatio-temporal neural activity was correlated with a given set of CNN features, we regressed the neural responses in the source space against the CNN-derived features of each image. Following obtaining the dSPM solutions for each image, an ordinary least square regression was run for each participant, for each time point and each source point. The coefficient of determination (or R-squared), indicating the proportion of variance explained by the regressors (CNN features) was used as the summarizing statistic to characterize the correlation between neural responses and a given set of CNN features.

### 2.8 Preprocessing of fMRI data and definition of regions of interest (ROIs)

The fMRI localizer data were preprocessed in SPM12 (http://www.fil.ion.ucl.ac.uk/spm/software/spm12/). The preprocessing included an unwarp transform to correct for geometric distortions using the fieldmap scan, a frame-by-frame transform to correct for head motion combined with a transform to align with the structural MRI, and finally spatial smoothing with an isotropic Gaussian kernel (where the full width at half maximum was 4 mm). The data in all localizer runs for each participant were concatenated, high pass filtered with a cut-off frequency at 0.0078125 Hz (a 128-second period), and then analyzed using a “general linear model” in a block design. In this model, the time series at each voxel was linearly regressed against a design matrix, which included, in separate columns, the pre-defined canonical hemodynamic response function convolved with the square-wave-like indicators of blocks for each stimulus condition (scenes, weak contextual objects, and phase scrambled scenes). The design matrix also included extra columns corresponding to nuisance covariates (e.g., the time series of parameters in motion correction). An autoregressive model of order 1 (AR(1)) was used to account for the temporal correlations in the residuals.

Scene/object selective regions were defined using the `MarsBaR` toolbox (http://marsbar.sourceforge.net/index.html). For any voxel, let *β*_scene_ denote the regression coefficient for the scene condition, *β*_object_ for the weak-contextual object condition, and finally *β*_scramble_ for the phase-scrambled scene condition. The *t*-statistics of the difference (*β*_scene_ − 1*/*2(*β*_object_ +*β*_scramble_)) was computed as the estimated difference divided by the estimated standard deviation of the difference. Then the voxels where the *t*-statistics was above a threshold were selected as scene-selective voxels. A customized threshold was set for each participant, such that clusters of contiguous voxels above the threshold were identified within or in the proximity of the parahippocampal gyrus, the retrosplenial cortex, and the transverse occipital sulcus in each hemisphere. These clusters were labeled as the scene-selective ROIs, which were the parahippocampal place area (PPA), the retrosplenial complex (RSC) and the occipital place area, also know as the transverse occipital sulcus (TOS). For the majority of the participants, the threshold was equal to the value where the family-wise error rate was controlled at 0.05. For some individuals, if the threshold was too stringent, we relaxed the threshold to a smaller value. Similarly, object-selective clusters in the lateral occipital cortex and the fusiform area (the lateral occipital complex or LOC) were also identified in both hemispheres for each participant respectively, where the difference of interest was *β*_object_ − *β*_scramble_.

The `SUMA` software (https://afni.nimh.nih.gov/Suma, Cox (1996)) was used to project these clusters of voxels to sets of vertices on the cortical surfaces, so that they could be labeled in the source space in MEG. The mapped sets of vertices were then manually examined, and the ones that had fuzzy boundaries or that were anatomically off were corrected. Each remaining contiguous set was defined as one region of interest (ROI). The ROIs that did not cover at least 10 source points (in MEG source space) were dilated until they covered 10 source points. Finally, the vertices in the LOC that were also in one of the scene-selective ROIs (PPA, RSC or TOS) were removed from the LOC. Additionally, some ROIs were defined based on the parcellation of the structural MRI by `Freesurfer`. These ROIs included the left and right pericalcarine areas that covered the early visual cortex (EVC). The pericalcarine areas included mostly V1 but might also include some part of V2. Finally, each pair of the corresponding bilateral regions were merged into one ROI, resulting in the bilateral PPA, RSC, TOS, LOC and EVC. See the supplementary materials for the locations of the ROIs for each participant.

### 2.9 Extracting features of the images

We used a convolutional neural network called Alexnet (Krizhevsky et al., 2012) implemented in the `Caffe` (Jia et al., 2014) software to extract sets of CNN features. This CNN was trained to classify images into 1,000 object categories with 1.2×10^6^ training samples and 5×10^4^ validation samples in the `ImageNet` database (Deng et al., 2009; Russakovsky et al., 2015). Figure 2 shows the 8-layer architecture of `Alexnet`. The first 5 layers had convolutional units. Each unit in these layers applied a dot product of a “kernel” weight matrix with the inputs within a receptive field—for example, in Layer 1, each unit’s receptive field was 11×11 pixels of the RGB channels of the raw image. Within each of the convolutional layers, there were a number of sub-layers; for example, Layer 1 had 48×2 = 96 sub-layers. Units in each sub-layer shared a “kernel” weight matrix, and therefore in such a convolutional architecture, the number of parameters was much smaller than that of an all-to-all connected neural network with the same number of units, yielding a more tractable model to train. After the convolution operation (dot product), each unit then applied a rectified linear function f(*x*) = max(0, *x*) on the dot product to generate the output. In Layers 1, 2, and 5, an additional normalization step was applied, where in each location, the output of the unit in each sub-layer unit was normalized by a function of the sum of squares of the responses in its neighbor sub-layers, including itself; a max-pooling operation was also added after the normalization in these layers (for details, see Krizhevsky et al. (2012)). Layers 6 and 7 were fully connected layers, each consisting of 2048×2 = 4096 units, and Layer 8 was the final output layer with 1,000 units, corresponding to the 1,000 object categories. We downloaded a pre-trained version of `Alexnet` from the `Caffe` “model zoo”, and re-sized our 600×600 images (including the gray box) to the input size required by `Alexnet`. For each image, we collected responses for all of the units in each layer and concatenated them into a vector. For Layers 1, 2 and 5, we used the responses before normalization and max-pooling.

**Figure 2:**
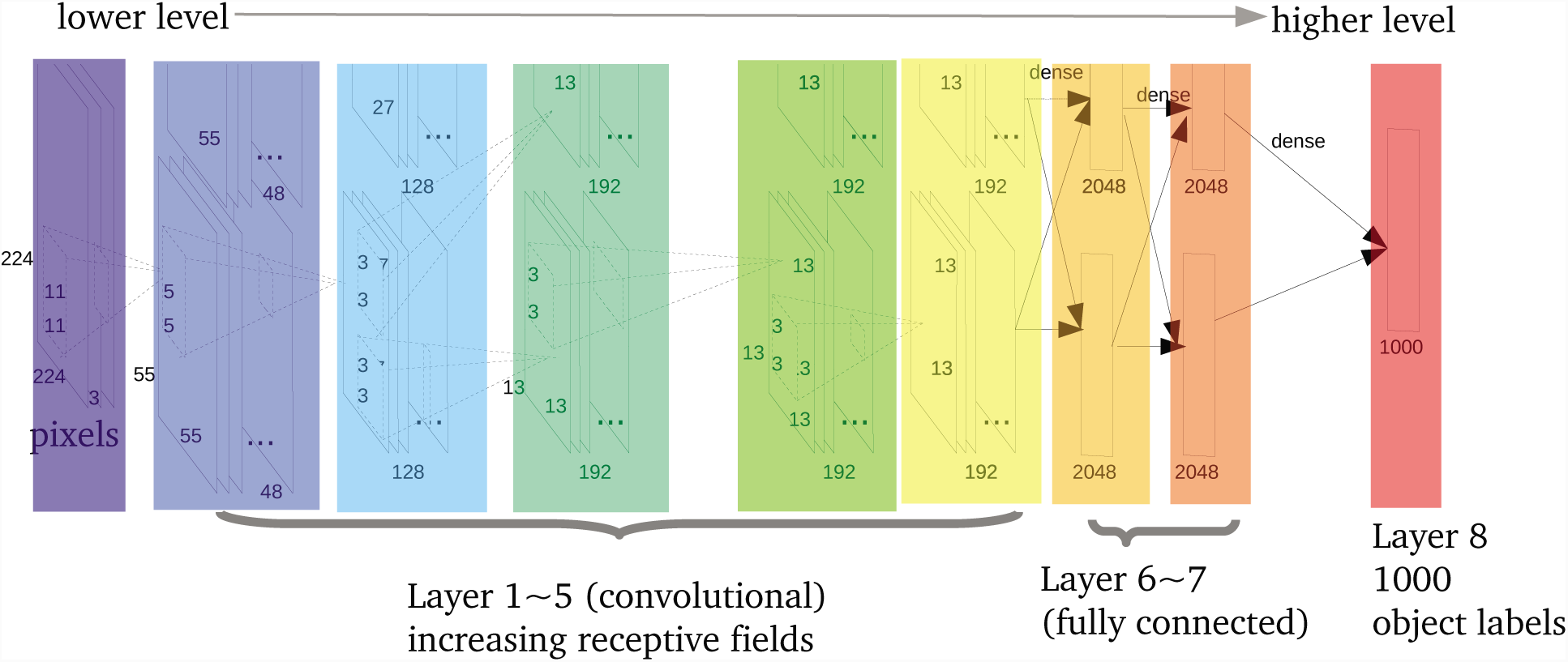
Structure of `Alexnet` (Krizhevsky et al., 2012). Note that the network was distributed on two graphics processing units (GPUs), illustrated by the upper and lower rows for each Layer. Extracted responses of the units were concatenations of the responses of all units in both the upper and lower rows.

Note that as with all CNNs, the 8-layer structure of `Alexnet` is feedforward; it naturally provides a progressive shift from low-level to high-level features. Here we chose Layer 1 and Layer 7 as representatives of low-level and high-level (object-category-related) features, respectively. Consistent with this approach, Layer 1 had the smallest receptive field sizes and the convolutional “kernel” weight matrices were similar to 2-D Gaussian functions and Gabor filters (see visualization in Krizhevsky et al. (2012)). In contrast, Layer 7 was the last fully-connected hidden layer before the output layer and was expected to represent task-relevant semantic information about the images. We did not include Layer 8 features. This decision was based on the observation that the correlation effect between the neural activity and Layer 8 features was much weaker than with other layers (see Figure 3 below). Layer 8 was the output layer, and the output 1,000 category labels were defined according to WordNet (Fellbaum, 1998; Deng et al., 2009), instead of a data-driven manner using brain activity. Therefore, these categories may not necessarily represent the organization of objects in the visual cortex.

**Figure 3:**
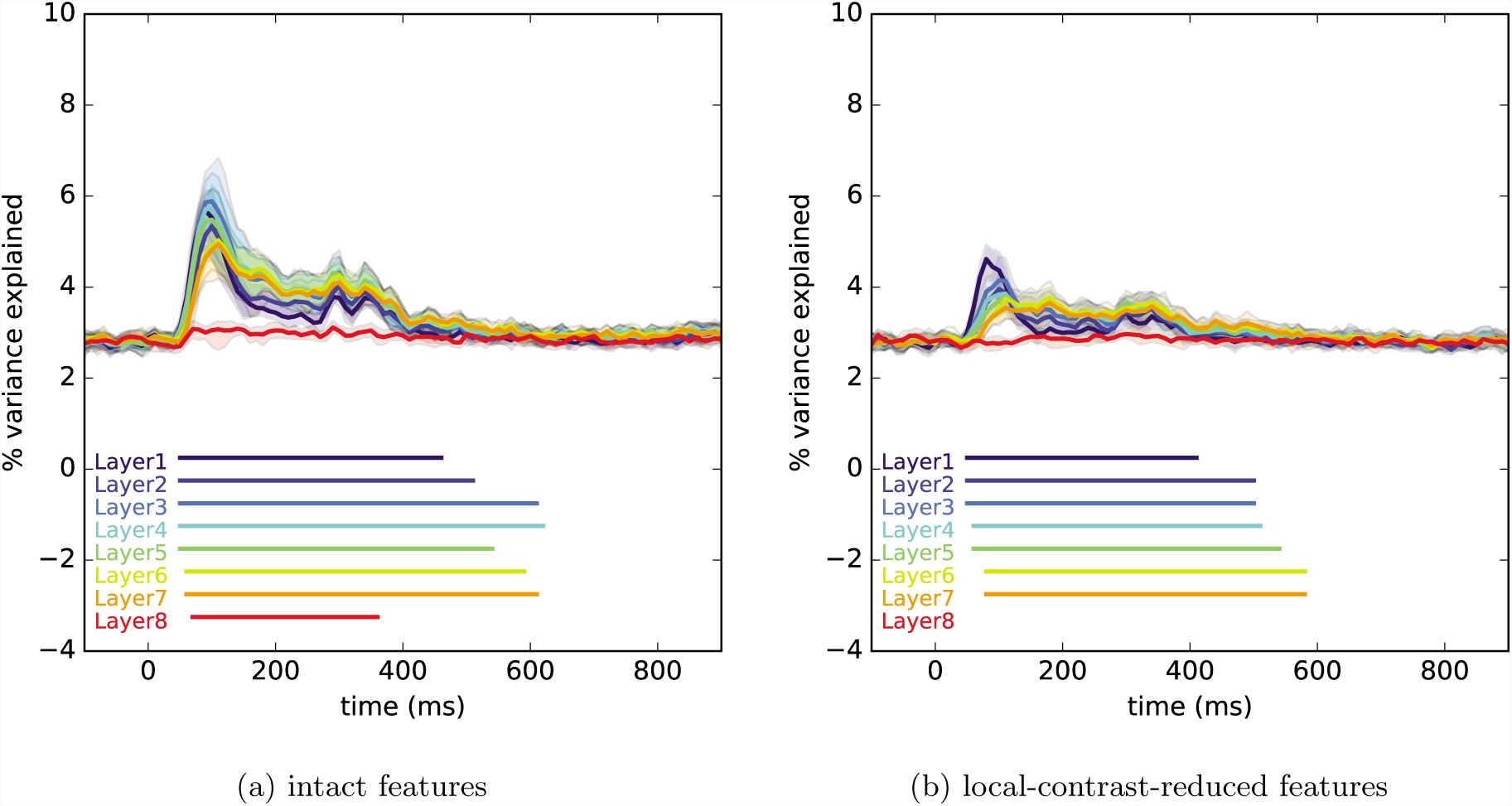
Proportion of variance explained by the first 10 principle components of the AlexNet features in each layer, averaged across sensors.

As mentioned earlier, nuisance covariates due to the various widths and heights of the stimulus images might also have been correlated with the extracted CNN features. Therefore we regressed out the width, height, area (width×hight) and aspect ratio (width */* height) from the features extracted from each unit across the union of the stimulus set and additional image set. An all-one column was included in the regression, which removed the mean across all images.

In addition to the `AlexNet` features, we also extracted simpler low-level features—the `local contrast` features—in the following way. For each image resized to the input size of Alexnet, the patch within each 11 × 11 receptive field in Layer 1 was converted to gray values (by averaging the values of the RGB channels), and the contrast in the patch was defined as (*x*_max_ − *x*_min_)*/*(*x*_max_ + *x*_min_), where *x*_min_ and *x*_max_ were the minimum and maximum of the gray values in the patch. The contrast values were concatenated across all receptive fields, yielding a 55×55 = 3025-dimensional vector for each image. Linear projections onto the nuisance covariates related to the widths and heights of the images were removed.

Note that the low-level and high-level layers in `AlexNet` are both highly correlated with `local contrast`, suggesting that there is some potential for local contrast alone to naturally elicit significant neural responses in the visual cortex. To account for this possible confounding effect due to `local contrast`, we regressed out the first 160 principal components of the `local contrast` features (explaining 90% of the variance in the local contrast features) from the intact features in all layers in `AlexNet`, and took the residuals as new features of interest. We term these new features `local-contrast-reduced features` hereafter.

### 2.10 Identifying common and residual feature sets

Within AlexNet, as one progresses from Layer 1 to Layer 7, the units in each layer non-linearly transform the raw pixel inputs to informative task-related features (where the task `AlexNet` was trained on was object categorization). In this context and as discussed earlier, Layer 1 features of `AlexNet` were identified as low-level features, while Layer 7 features were identified as high-level features. However, this does not mean the features in Layer 1 and Layer 7 are completely orthogonal to one another. More specifically, there is likely to be some linear dependence between the low-level Layer 1 feature set and the higher-level Layer 7 feature set. As such, correlating neural data separately with both Layer 1 and Layer 7 does not allow us to infer the degree to which these different correlations are due to the common linear components between the two layers. To address this ambiguity, we used canonical correlation analysis (CCA) to extract the *common linear components* between Layer 1 and Layer 7. In particular, for two multivariate variables, canonical correlation analysis learns a linear projection for each variable, such that the correlation between the projected results from the two variables is maximized. Similar to principal component analysis, we can learn more than one pair of linear projections, thereby obtaining correlated components that are orthogonal with each other.

Because the CNN features were high dimensional, we included a set of additional images to boost our ability of analyzing the features. These additional images were from the same 181 scene categories in the same image dataset, including 6 exemplars per category, different from the stimulus images presented in the MEG experiment (6×181 = 1086 images in total); they had similar sizes (longest side = 500 pixels) as the stimulus images, and were also centered in the same 600×600 bounding boxes. The CNN features of these images were obtained in the same way as the stimulus images. The local contrast features for each additional image were obtained as well; across both stimulus images and additional images, these local contrast features were used in the principal component analysis (PCA) to obtain the 160 principal components, and, these 160 principal components were regressed out form the features in each layer across all images, resulting in the `local-contrast-reduced features`, which were used for the analysis below.

Both Layer 1 and Layer 7 features are high-dimensional (290400 for Layer 1 and 4096 for Layer 7); in contrast, there were only 362 stimulus images. To reduce dimensionality of the feature sets for the CCA analysis, we ran principal component analysis (PCA) on the Layer 1 and Layer 7 features of the union of the 362 stimulus images and the 1086 additional images. We then retained only the first 362 principal components. We extracted the CCA linear projections with *p*_*c*_ = 6 components using only the dimension-reduced features for the 1086 additional images. The projections were then applied to the dimension-reduced features corresponding to the 362 stimulus images; following this projection, we have 362×6 new features from Layer 1 and 362×6 new features from Layer 7, where the correlation between corresponding dimensions ranged from 0.8 to 0.5. These two sets of features were concatenated into a 362×12 matrix and a further PCA was used to just pick the first 6-dimensional components (in that every corresponding pair was highly correlated, 6 components were able to explain the bulk of the 12 dimensions). We term this new 362×6 feature matrix the *common components*.

The choice of *p*_*c*_ was based on a 6-fold cross-validation analysis using only the additional images. For *p*_*c*_ = 1, *…,* 20, we used 5 fold of the additional images to learn a CCA model, and then fit a linear mapping between the projected components in Layer 1 and Layer 7, whose correlation was already maximized. On the last fold of testing data, we predict the Layer 1 features using the Layer 7 features, via the trained CCA projections and the linear mapping. We repeated the same procedure to predict Layer 7 features using Layer 1 features as well. We chose the maximum *p*_*c*_ where the cross-validation error was below chance for at least one direction of prediction — the two dictions being Layer 7 to Layer 1 and Layer 1 to Layer 7. This resulted in *p*_*c*_ = 6. The cross validation error increased substantially after *p*_*c*_ *>* 6.

After obtaining the *common components*, we computed a feature set we will refer to as *residual Layer 1* of the 362 stimulus images by regressing out the *common components* from the dimension-reduced Layer 1 features and then picking the first 6 principal components of the residuals. A similar feature set, which we will refer to as *residual Layer 7*, was computed for Layer 7 in the same way. Note that *common components, residual Layer 1*, and *residual Layer 7* were all 6-dimensional; therefore we can compare the neural correlation profiles for each of these three feature sets. Mathematical details and the cross-validation results can be found in Chapter 5.2.10 and 5.3.1 in Yang (2017).

Given how *common components, residual Layer 1*, and *residual Layer 7* were created, we suggest the following intuitive interpretation. The *common components* are the linearly correlated components between the low-level Layer 1 features and the high-level Layer 7 features. Since the features in Layer 7 are highly relevant to the object-category labels, *common components* represents the “object-category-relevant” or task-relevant low-level features, for example, informative edges that defines the shape of an object. In contrast, *residual Layer 1* represents components in Layer 1 features that were roughly orthogonal to higher-level semantic information that is relevant to the task of object categorization. Finally, *residual Layer 7* represents linear components in Layer 7 that were roughly orthogonal to features in Layer 1; in other words, *residual Layer 7* represents unique high-level features beyond the information captured in the low-level features.

### 2.11 Confidence intervals and statistical tests

#### Percentile confidence intervals

After obtaining the statistics representing the regression effects (e.g., R-squared) for each time point, we computed the group-averaged time series across participants, for which the confidence intervals were obtained through bootstrapping. We randomly re-sampled the time series of statistics at the participant level with replacement, and used the (*α*_0_, 1*-α*_0_*/*2) percentile confidence intervals of the bootstrapped sample (Wasserman, 2010). The significance level *α*_0_ here was defined as 0.05*/T/n*_*stat*_, where *T* was the number of time points in the time series, and *n*_*stat*_ was the number of time series of statistics that were considered (Bonferroni correction). For example, in cases where we plot the R-squared for 8 layers of features, *n*_*stat*_ = 8.

#### Permutation-excursion tests

When examining whether a time series of statistics is significantly different from the null hypothesis in some time windows, it is necessary to correct for multiple comparisons across different time points. Here, permutation-excursion tests (Maris and Oostenveld, 2007; Xu et al., 2011), were used to control the family-wise error rate and obtain a global *p*-value for across time windows. In a one-sided test that examines whether some statistics were significantly larger than the null, we first identified clusters of continuous time points where the statistics were above a threshold, and then took the sum within each of these clusters. Similarly, in each permutation, the statistics of permuted data were thresholded, and summed within each of the detected clusters. The global *p*-value for a cluster in the original, non-permuted case was then defined as the proportion of permutations where the largest summed statistics among all of the detected clusters was greater than the summed statistics in the cluster from the non-permuted case.

To test whether the mean time series of some statistics (e.g., R-squared in regression) across participants was significantly larger than that in the baseline time window before the stimulus onset, we computed the difference, separately for each participant, between the original time series and the temporal mean of the statistics within the baseline time window. Then across participants at each time point, we used the *t*-statistics defined in the Student’s *t*-tests to examine if the group means of these differences were significantly above zero in any time window. Each permutation was implemented by assigning a random sign to the difference time series for each participant. This test, which we refer to as a *permutation-excursion t-test* hereafter, was implemented in `MNE-python`, where the number of permutations was set to 1024. The threshold of the *t*-statistics was equivalent to an uncorrected *p*-value ≤0.05.

#### Other corrections of multiple comparisons

In addition to the permutation-excursion tests discussed above, in cases where the possible dependence structure was less readily characterized as compared to that in adjacent time points, we relied on other methods to correct for multiple comparisons and control for the false discovery rate, including the Benjamini-Hochberg procedure (Benjamini and Hochberg, 1995).

## 3 Results

By regressing neural activity at different time points and brain areas onto stimulus features extracted from a pre-trained CNN, we obtained spatio-temporal correlation profiles that offer insights as to information flow in the human vision cortex. Below we present these correlation profiles in both MEG sensor space and source space as defined on the cortical surface. As an overview, we first examined how each layer of `AlexNet` accounted for variance of neural activity. Next, we decomposed low-level and high-level AlexNet features into three orthogonal sets: the common components between low-level and high-level features, the residual low-level features and the residual high-level features; we present the neural correlation profiles with these sets and analyze how these profiles not only support feedforward information flow but also indicate non-feedforward information flow when participants process naturalistic scene images.

### 3.1 Sensor-space regression

To examine whether the features extracted from different layers in the pre-trained CNN (AlexNet) are able to explain the MEG data in sensor space, we ran an ordinary least square regression analysis at each sensor and each time point for each participant. This approach allowed us to compare the linear dependence (or correlation) between different feature sets and neural activity at different time points. For this regression, the neural responses to all 362 images were included.

Because `AlexNet` features are high-dimensional as compared to the number of observations, we need to avoid overfitting through either dimensionality reduction or other regularization methods. In that our main goal was to test whether a significant amount of variance was explained by each layer and for computational simplicity, we used principal component analysis (PCA) to reduce the dimensionality and included the first 10 principal components as regressors. The choice of 10 was admittedly arbitrary; we expect similar results so long as the number of components is neither too small (e.g., 2-3) or too large such that the regression models overfit. We used the first 10 principal components of both the intact features and the *local-contrast-reduced features* from each layer as regressors. For the *local-contrast-reduced features*, the variation capturing contrast in local neighborhoods at different locations of an image was removed (see Materials and methods).

We quantified the correlation between neural activity and the regressors in each layer as the proportion of variance explained (i.e., R-squared). In other words, R-squared is the statistic in our correlation profiles. For purposes of visualizing overall effects, R-squared was averaged across all sensors at each time point for each participant. Figure 3 illustrates these results, where each curve represents the R-squared for each layer, averaged across all sensors and all participants. The transparent bands show the confidence intervals obtained by bootstrapping the observed R-squared time series at the participant level. *Permutation-excursion t-tests* were used to test whether the averaged R-squared across sensors was greater than the temporal average of that in the baseline time window (−140 to −40 ms with regard to the stimulus onset), during which MEG signals should be independent of the stimulus images, and thus the regressors. The significant time windows were identified where the *p*-values of the *permutation-excursion t-tests* were smaller than 0.05/8. These significant windows are marked by the colored segments under the curves in Figure 3. Note that these *p*-values were already corrected for multiple comparisons at different time points: the denominator 8 was used as a correction for the 8 tests corresponding to the 8 layers according to the Bonferroni criterion.

In Figure 3, the left plot shows the results from using the intact features, while the right plot shows the results from using the *local-contrast-reduced features*. In both plots, we identified significant time windows (from 80 ms to about 500 ms) for Layer 1 through Layer 7. In these windows, the variance explained by the features was significantly greater than that in the baseline time windows. These results indicate that the neural responses recorded by MEG in these time windows were correlated with the `AlexNet` features.

Because we used the same number of orthogonal principal components for each layer, the model complexity of the regression for each layer is the same. Thus it is fair to compare the R-squared statistics between layers and analyze which layer explained the neural responses better. Both plots show a pattern of early-to-late, lower-level to higher-level shift, where lower-level layers, especially Layer 1 and 2, explain a larger proportion of variance before 150 ms, and higher-level layers, especially Layer 6 and 7, explain a larger proportion of variance from about 150 to at least 400 ms. This pattern is consistent with feedforward information flow as assumed in hierarchical models of visual cortex. Interestingly, Layer 8 generally explains lower variance as compared to the other seven layers—no significant time window was detected in Figure 3(b). One possibility is that the object categories represented in Layer 8 did not align with the basic, neurally-discriminable categories that humans naturally use, or that the neural activity associated with object category labels is weaker than the neural activity associated with visual features.

Also of interest, although the proportion of variance explained by the principal components from the intact features appeared higher than the proportion of variance explained by the principal components from the *local-contrast-reduced features*, the early-to-late, lower-level to higher-level temporal patterns were much clearer in the latter case. These results indicate that using the *local-contrast-reduced features* may facilitate better differentiation in the neural correlations with feature sets at different levels. As such, throughout the remainder of this paper, we present only results with the *local-contrast-reduced features*; that is, unless there is specific note, “features” will refer to the *local-contrast-reduced features*.

The results discussed to this point only address temporal patterns. Next, we illustrate spatially which sensors demonstrated strong correlations with `AlexNet` features by visualizing the regression results on a topological map of the sensor layout. For each sensor, and for each time point from 50 ms to 700 ms with a 50 ms step-size, we averaged the R-squared over a 70-ms window centered at the time point and then across all participants. Note that the same sensor could map to different locations for different participants, due to individual variation in head sizes and the head locations in the MEG helmet. As a visualization, these results here are purely qualitative and primarily aid in a better understanding of overall patterns. We defer more rigorous statistical comparisons of the spatio-temporal correlation profiles to the source-space regression analysis.

Figure 4 includes the topological heat maps of the averaged R-squared (proportion of variance explained) for each MEG sensor by the first 10 principal components of the features in Layers 1, 3, 5 and 7. The strongest correlation effects were in the posterior sensors, which are physically close to visual cortex. From Layer 1 to Layer 7, we observe a shift of the correlation effects from early windows to late windows, especially within 50 ms to 250 ms time window. For Layer 1 (and also Layer 3), it is interesting that after the first transient peak at 100 ms, there appears to be a second peak from about 300 ms to 350 ms. Because each stimulus image was presented for 200 ms, we posit that this second peak may arise as a neural response to the disappearance of each image at 200 ms—there was roughly a 80 ms to 100 ms delay from the stimulus onset to the first peak, so we might expect a similar delay in the response to the disappearance of each image.

**Figure 4:**
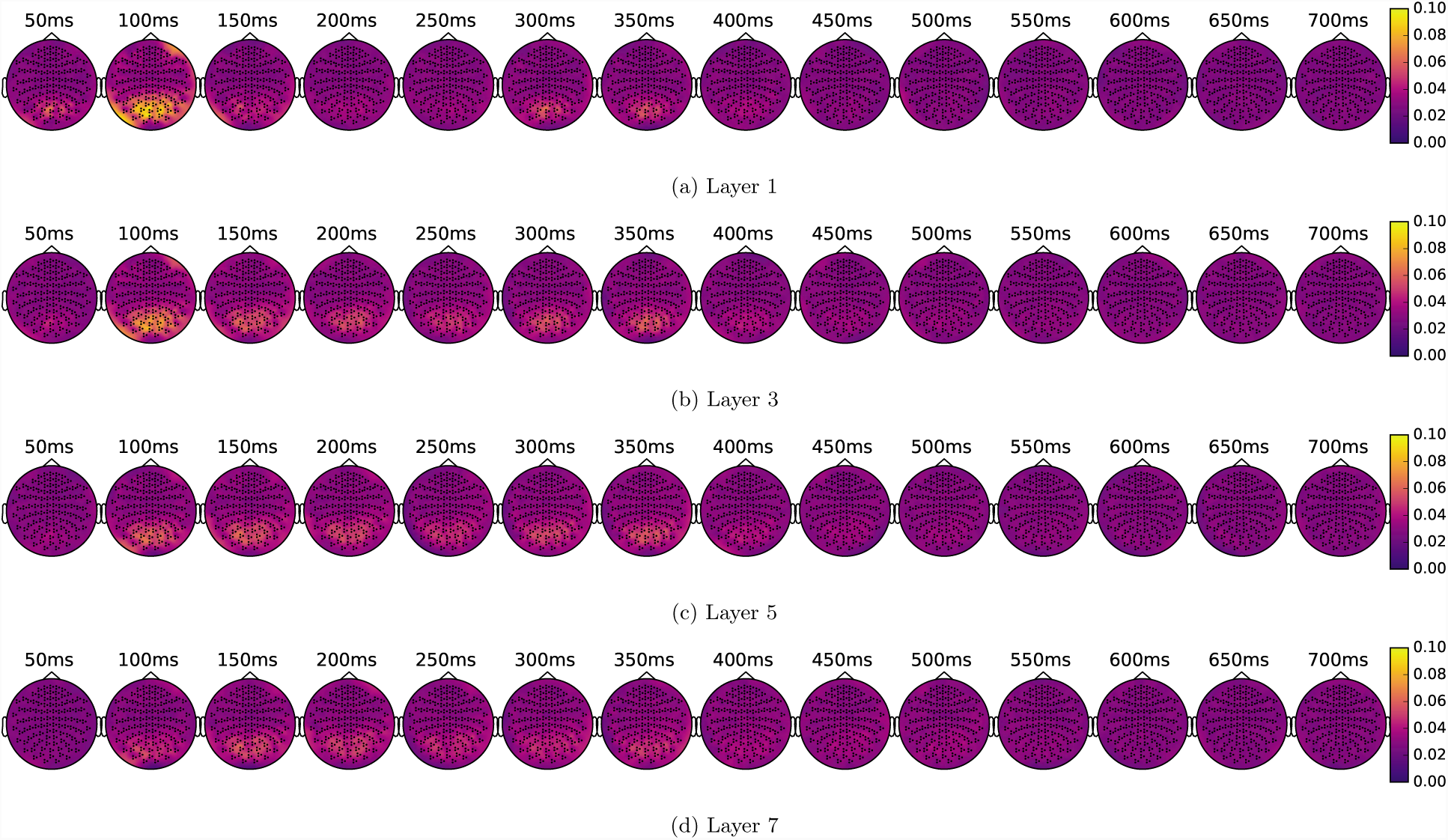
Topology maps of the proportion of variance (%) explained by the features from Layer 1, 3, 5 and 7.

Next, using Layer 1 as low-level features and Layer 7 as high-level features, we further derived three orthogonal sets of features from these starting points, using the canonical correlation analysis of the *localcontrast-reduced features* (*p*_*c*_ = 6, see Materials and methods): *common components* represents the shared linear components between Layer 1 and Layer 7, or the components in the low-level Layer 1 that are highly correlated with high-level features; *residual Layer 1* represents low-level features that are roughly orthogonal to Layer 7; *residual Layer 7* represents a high-level features that are roughly orthogonal to Layer 1. In Figure 5, we present the topological heat maps of correlation effects for these three sets of derived features.

**Figure 5:**
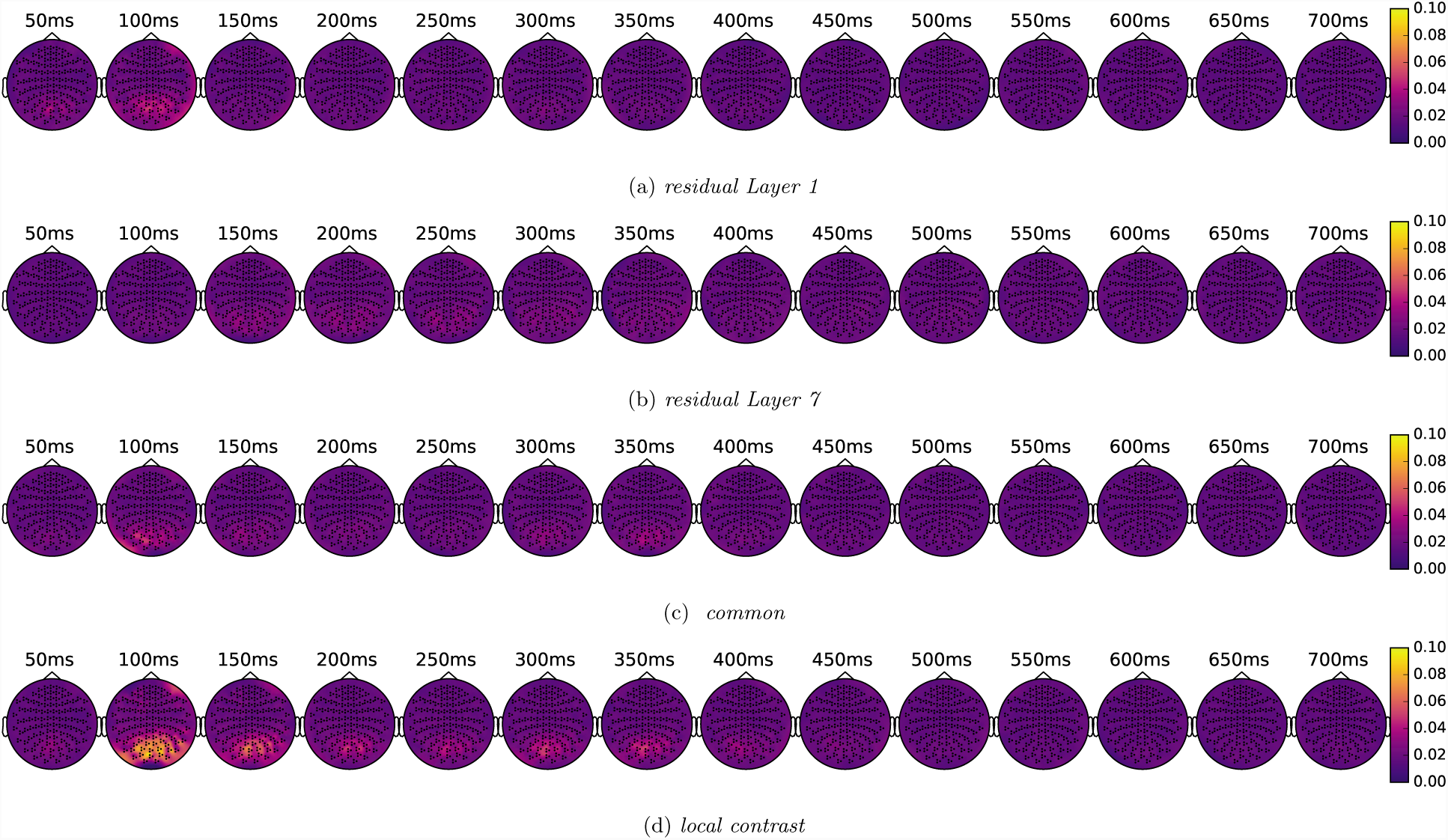
Topology maps of the proportion of variance explained by the *common components*, the *residual Layer 1, residual Layer 7* and the *local contrast features*.

As illustrated in Figure 5, the *residual Layer 1* exhibited an early peak near 100 ms, while the *residual Layer 7* exhibited correlation effects that were somewhat later, from about 150 ms to 400 ms. In contrast, the *common components* appeared to have two peaks, one centered at 100 ms, and one from about 300 ms to 350 ms. This latter peak may correspond to neural responses to the disappearance of each stimulus at 200 ms; nonetheless, there was not a significant later peak for *residual Layer 1*, near 300 to 350 ms, indicating that in this time window, the *common components* (low-level components that are correlated with high-level features) are represented differently from the *residual Layer 1* (the low-level components that are roughly orthogonal to high-level features). Comparing *common components* and *residual Layer 1* (both are low-level features) with the high-level feature set (*residual Layer 7*), we can again observe the early-to-late, lower-level to higher-level shift, consistent with feedforward information flow.

In the last row of Figure 5, for comparison, we also plot the topological heat maps of the correlation effects for the first 6 principal components of the *local contrast features*. Note that the local contrast features were orthogonal to the three sets of features above (see Materials and methods). The local contrast features showed large correlation effects spanning from 100 ms to 400 ms. These correlation effects were larger than those for the three sets above, indicating that local contrast did elicit relatively strong neural responses.

Summarizing the regression results in sensor space, we observed an early-to-late, lower-level to higher-level shift in the temporal correlation patterns, indicating a feedfoward information flow during visual scene perception. When we decomposed the *local-contrast-reduced features* from Layers 1 and 7 into three roughly orthogonal feature sets—the *common components*, the *residual Layer 1* and *residual Layer 7* —we observed an apparent temporal separation of the low-level and high-level features (e.g., when comparing the *residual Layer 1* and the *residual Layer 7* or comparing the *common components* and the *residual Layer 7*). In addition, the *local contrast features* explained a larger proportion of variance in the neural data than the three groups derived from the *local-contrast-reduced features*; suggesting that failing to partial out local contrast features from AlexNet features would result in the majority of observable correlation effects being due to the local contrast features.

### 3.2 Source-space regression

As presented above, the regression results in sensor space are only informative regarding temporal correlation profiles between neural activity and stimulus features. Here, we shift to source space to examine the spatio-temporal correlation profiles. Source localization was achieved using the dynamic statistical parametric mapping (dSPM) (Dale et al., 2000) method; in this way, neural activity was obtained at each source point in brain space as defined on the cortical surface. We then ran ordinary least regression analyses to obtain the correlation profiles. Note that source localization *per se* is a challenging problem, as the number of source points in brain space (about 8000 in our case) is an order of magnitude larger than the number of sensors (306). The dSPM method uses an *L*_2_ regularization, which penalizes the *L*_2_ norms of the source activity to obtain a unique solution. This is equivalent to placing a prior on the source neural activity because we have no further information about where to localize the signals. To verify that our results are not due to this specific source localization prior, in Supplementary Materials, we also applied an *L*_1_*/L*_2_-based sparsity-inducing source regression method, a short-time Fourier transform regression model (STFT-R, Yang et al. (2014)), and obtained qualitatively similar results.

Subsequent to source localization and regression, we mapped the source space of each participant onto a default template in `Freesurfer`. We then obtained whole-brain maps of the averaged R-squared statistics for the first 10 principal components for each layer in `AlexNet` (the same as in “Sensor-space regression”). Figure 6 shows visualizations at different time points from a ventral view of the cortical surface for Layers 1, 3, 5 and 7.

**Figure 6:**
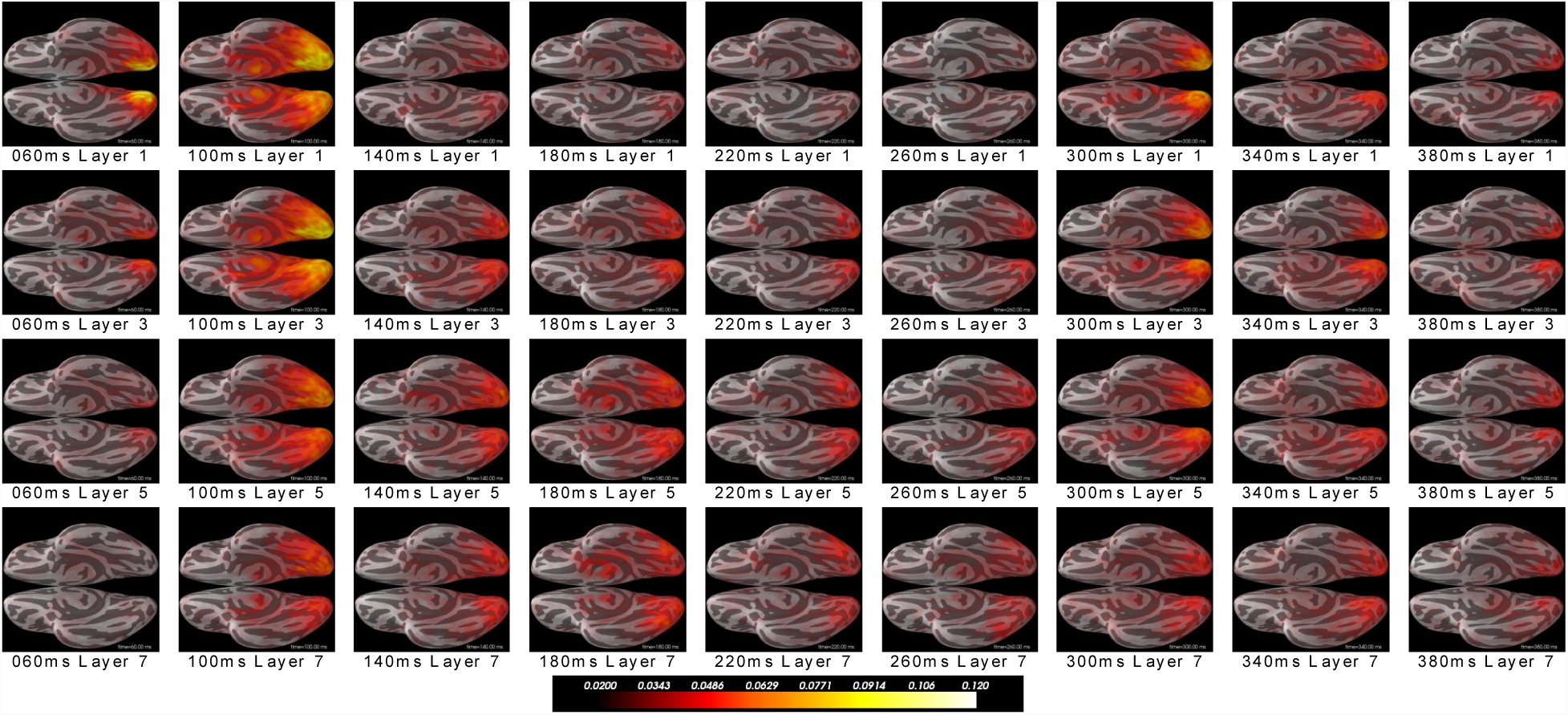
The proportion of variance explained by different layers, computed from the dSPM source solutions that were morphed onto a common template and aver1aged across participants.

Adjacent source points typically contribute to the sensor recordings in a similar way; as a result, *L*_2_-norm regularized methods such as dSPM generally have a spatial blurring effect, where an underlying single large current dipole can be reconstructed as distributed current dipoles covering a large cortical area. As a consequence, when we observe strong effects (i.e., large averaged R-squared) in large areas, this pattern may be due to either local large effects or genuinely distributed effects. Keeping this issue in mind, we examine our results in Figure 6. At 60 ms, the correlation effects localized in the posterior end near the early visual cortex were strongest for Layer 1, and as we move from Layers 3, 5 and 7, the regression effects gradually decrease: at 100 ms, the regression effects had large magnitudes and spread over large areas of the ventral visual cortex; at 140 ms to 180 ms, the correlation effects appeared stronger and in larger areas for Layers 3, 5 and 7 than for Layer 1, and the effects spread to more anterior regions for Layers 5 and 7. This overall pattern is consistent with feedforward information flow along the hierarchy from posterior to anterior regions of visual cortex. In addition, at approximately 300 ms to 350 ms, we observe a second correlation peak for most layers in posterior regions close to early visual cortex. This later onset peak shows a delay of roughly 100 ms from the disappearance of each stimulus at 200 ms—similar to the delay of the first peak from the stimulus onset—as before, this later peak may be a result of a correlation between neural responses to the disappearance of the images and the CNN features of the images.

Next, we present the results of the source-space regression analysis using the three sets of derived features, the *common components*, the *residual Layer 1* and the *residual Layer 7*. After obtaining the dSPM source estimates, we ran a linear regression for each source point at each time point, using each of the three feature sets as the regressors to obtain the R-squared statistics. Each time series of R-squared values may potentially have different magnitudes across different source points and participants, so we further normalized these time series using the following method. For each source point, we divided the R-squared values at each time point by the sum across all time points and all three groups. In order to conduct rigorous statistical tests to compare the correlation profiles between these three feature sets, we focused on several representative ROIs along the visual hierarchy, including the pericalcarine areas that covered the EVC, and the object/scene-selective areas (LOC, PPA, RSC and TOS). This ROI approach was in contrast to a whole-brain analysis, where we would have needed to control for multiple comparisons at thousands of source points. In particular, by aggregating the correlation effects within each ROI, we had fewer multiple comparisons and therefore higher statistical power. Since these regions were defined individually for each participant, we also did not have to morph the individual source spaces onto a template. Our analysis proceeded by averaging these normalized R-squared values across the source points within each ROI. Note that these normalized values reflect the relative correlation effects between localized neural activity and the three groups of regressors within the ROI, and thus they represent the spatio-temporal profiles of the correlation effects we will focus on hereafter.

Figure 7 shows the correlation profiles and related comparisons within each ROI. The first column shows the average of the normalized R-squared values across participants for each group of regressors within each ROI. The transparent bands show 95% confidence intervals, bootstrapped at the participant level and corrected for multiple comparisons across all time points and for the three regressor groups using the Bonferroni criterion. We also ran pairwise comparisons between the correlation profiles of the three feature sets. In these pairwise comparisons, for each source point, each time point, and each participant, we computed the ratio of the R-squared value for each regressor set (i.e., each feature set) to the sum of the R-squared values across the three groups. In this way, each time point had comparable statistics that described the relative strength of the correlation effects for each of the three regressor groups. Again, for each participant, we averaged these ratios across the source points within each ROI. We then took pairwise differences (*residual Layer 7-residual Layer 1, residual Layer 1-common components*, and *residual Layer 7-common components*), and examined whether the averaged differences across participants were significantly different from zero in each ROI. The remaining three columns in Figure 7 show the averaged differences between each pair of the three sets of features: the second column (cyan) corresponds to the difference between *residual Layer 7* and *residual Layer 1*; the third column (yellow) corresponds to the difference between *residual Layer 1* and *common components*; the fourth column (magenta) corresponds to the difference between *residual Layer 7* and *common components*. The transparent bands show the bootstrapped 95% confidence intervals, corrected for the three comparisons at all the time points using the Bonferroni criterion. *Permutation-excursion t-tests* were used to identify time windows where the two-sided *p*-values were smaller than 0.05. Note that in this case, we only corrected for multiple comparisons across time points. To further correct for comparisons across different pairs and multiple ROIs, we controlled the false discovery rate (FDR) at 0.05 using the Benjamini-Hochberg procedure. The gray boxes in Figure 7 indicate the time windows that survived the correction (i.e., the difference between the correlation effects for the two sets of features was significantly greater or smaller than zero), and the *p*-values of the *permutation-excursion t-tests* (before the FDR correction) are marked. Note that there appeared to be some time windows that did not survive the correction, but in which we could visually observe some possibly non-zero differences based on the confidence intervals (which were not corrected for multiple ROIs). However, we were not able to claim that these windows had significantly non-zero differences in the comparisons. Nevertheless, even if the differences at some time points were not significant, there may still exist true underlying differences that we are unable to detect given the statistical power of our study.

**Figure 7:**
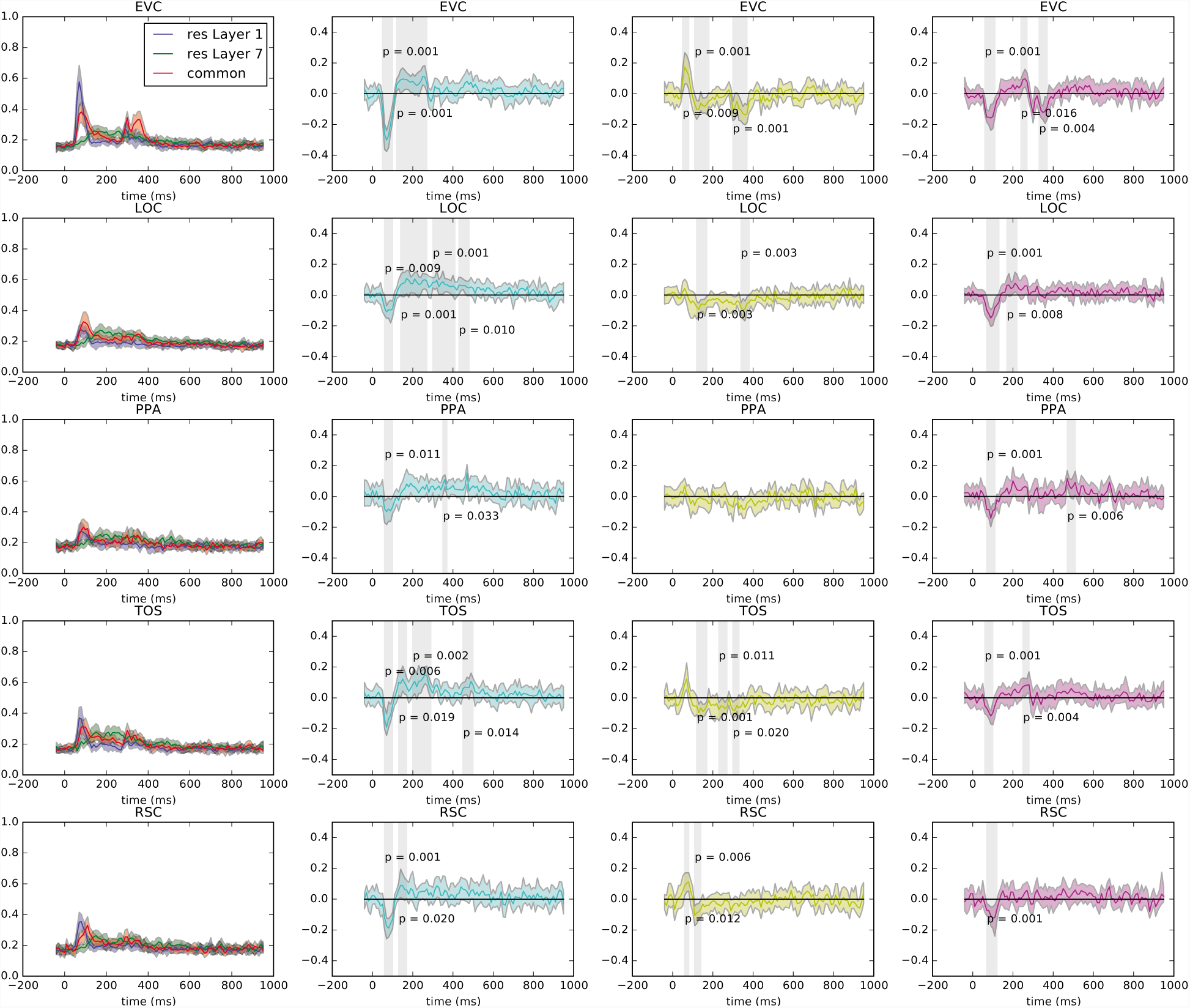
Correlation profiles in the ROIs in the source space. First column: averaged statistics of the correlation effects across participants (color code: blue: *residual Layer 1*; green: *residual Layer 7*; red: *common components*). Second to fourth columns, pairwise differences among the three feature groups (color code: cyan: *residual Layer 7-residual Layer 1*; yellow: *residual Layer 1-common components*; magenta, *residual Layer 7-common components*). The transparent bands indicates bootstrapped confidence interval at the participant level. Gray areas indicate time windows where the differences were significantly non-zero.

Next we analyze the correlation profiles and the pairwise comparisons for each ROI. In addition to contrasting the patterns between the three groups within each region, we also focused on the temporal changes of the correlation effects. In addition, we qualitatively compared correlation profiles in early visual cortex (EVC) and in object/scene selective regions (e.g., LOC, PPA) that are at higher levels than the EVC.

In the EVC (the first row and the first column, or see Figure 8 for an enlarged view), the *residual Layer 1* and the *common components* (the blue and red curves respectively)—corresponding to the low-level features roughly orthogonal to high-level features, and the low-level features that were correlated with high-level features (i.e., object-category-relevant)—had early transient correlation effects within 60 ms to 120 ms, peaking near 100 ms. The correlation effects of the *residual Layer 7* (the green curve)—corresponding to the high-level features that were roughly orthogonal to low-level features—increased later, starting at about 100 ms, peaking near 140 ms, and lasting until at least 400 ms. From the pairwise comparison between *residual Layer 7* and *residual Layer 1* (the cyan curve in the second column, the first row of Figure 7), we can see a negative peak within 60 ms to 100 ms, indicating that the correlation effect of the *residual Layer 7* was smaller than that of the *residual Layer 1*. Following this time window, the difference became positive from about 120 ms to 260 ms, indicating that the correlation effect of the *residual Layer 7* was greater than that of the *residual Layer 1*. These results are consistent with the patterns of the blue and green curves shown in the first column, which reveal an early-to-late shift of the correlation effects from low-level to high-level features.

**Figure 8:**
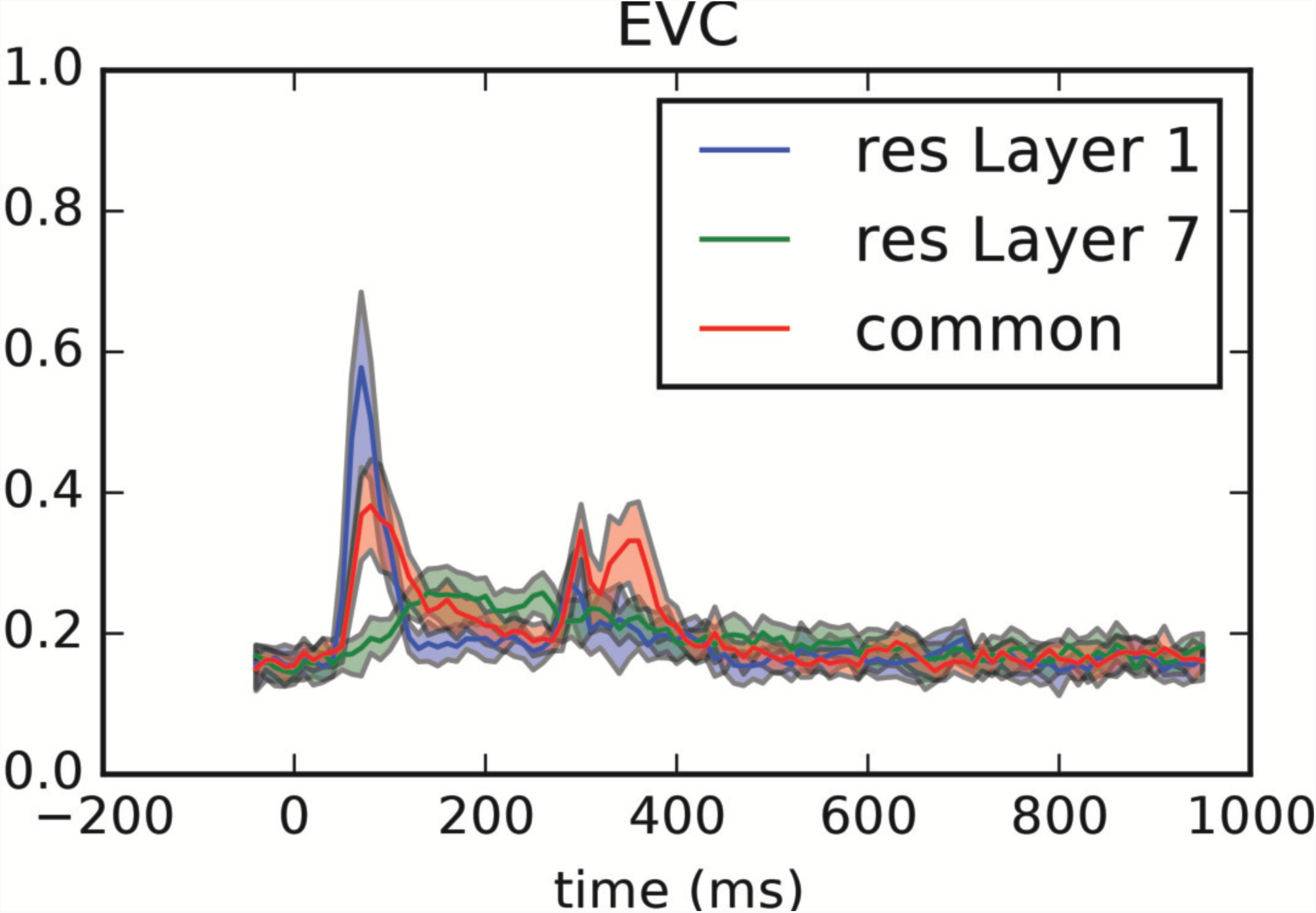
Correlation profiles in the EVC in the source space (enlarged from Figure 7)

Notably, the *residual Layer 1* and the *common components* both exhibited a second peak near 300 ms (in addition to the peak at 60 ms to 120 ms). However, this second peak appeared smaller and more transient for the *residual Layer 1* than for the *common components*; the correlation effect also lasted longer (until at least 380 ms) for the *common components* (the red curve) than for the *residual Layer 1* (the blue curve). This can be verified with the significant negative window in the pairwise comparison between *residual Layer,1* and *common components* (the first row, the third column of Figure 7, yellow), near 300 to 380 ms. In contrast, around the first transient peaks at 60 ms to 120 ms for *residual Layer 1* and *common components*, the difference (*residual Layer 1-common components*) was positive at the beginning and negative later, indicating that the correlation effect with *residual Layer 1* was higher, but that the correlation effect with the *common components* lasted slightly longer than the *common components* in the falling phase of the blue and red peaks as shown in the first column of Figure 7. In the pairwise comparison between the *residual Layer 7* and the *common components* (the first row and the the fourth column Figure 7, magenta), we observed negative peaks from about 60 ms to 120 ms and 350 ms, corresponding to the two transient red peaks of the *common components* in the first column. There are some significant positive differences between the time windows of these two transient peaks, indicating higher correlation effects with the high-level features (*residual Layer 7*) as compared to the object-category-relevant low-level features (*common components*).

One of the key findings of this study is the different correlation effects observed for the EVC with respect to *residual Layer 1* and *common components* starting at about 300 ms. This result indicates that the EVC is able to differentiate object-category-relevant and less-object-category-relevant low-level features. The specific timing of the these peaks at 300 ms may be due to the disappearance of the stimuli at 200 ms (i.e., EVC was sensitive to changes of visual inputs that were related to the low-level image features that disappeared). Nevertheless, compared with the first peaks of the blue and red curves near 60 to 120 ms (Figure 8)—due to the initial responses to the stimulus onset—the dynamics of these late transient peaks was dramatically different. In particular, in the early peaks, the correlation effect with the *residual Layer 1* (blue) was larger and the correlation effect with the *common components* (red) was just slightly longer than the blue peak; whereas in this later time window, the red peak was higher and lasted much longer. If the late red and blue peaks starting at 300 ms are due to the disappearance of the stimuli, some non-feedforward dynamics, either due to top-down feedback or local lateral connections appears to have “guided” the EVC to focus more strongly and for longer on object-category-relevant low-level features.

In an objective selective region, LOC (second row of Figure 7), which is at higher level than EVC along the hierarchy, the early transient peaks of the regression effects for the low-level features (the *residual Layer 1* and the *common components*) were much smaller, whereas the magnitude of the later peak for the high-level features (*residual Layer 7*) was similar to that in the EVC. Moreover, we did not observe second peaks for the *residual Layer 1* and the *common components* that were as prominent as those observed in the EVC. In the LOC, the pairwise difference plotted in the second column of Figure 7, (cyan, the second row, *residual Layer 7-residual Layer 1*) shows a similar pattern to that in the EVC, but the early negative difference was smaller and the later positive difference lasted longer, corresponding to the profiles of the correlation effects in the first column of Figure 7. In the pairwise difference plotted in the third column Figure 7, (yellow, the third row, *residual Layer 1-common components*), there was both an early and a late negative windows, indicating that the correlation effect of the *common components* was larger than that of the *residual Layer 1*. These results are consistent with a hierarchical organization for visual cortex, in that the LOC, being at a higher level in the hierarchy, showed higher correlation effects with the object-category-relevant low-level features (*common components*) as compared to the low-level features that were roughly orthogonal to high-level features (*residual Layer 1*). In the pairwise difference plotted in the fourth column of Figure 7, (magenta, the fourth row, *residual Layer 7-common components*), we observe an early negative window and a later positive window, indicating a temporal shift from higher correlation with the object-category-relevant low-level features to higher correlation with the high-level features, again consistent with feedforward information flow.

One might have expected that the LOC, which is at a higher-level within the hierarchy than the EVC, would show smaller correlation effects with low-level features than we actually observed. We speculate that because the columns in the forward matrix inherently have strong spatial correlations, the reconstructed source solutions can be spatially blurred, and the correlation effects with low-level features can “leak” from lower-level visual areas into the LOC. Moreover, even in the absence of spatial blurring, the hierarchy can be gradual such that we may only observe small relative differences between the correlation profiles between the EVC and the LOC.

The third row Figure 7 shows the results for a scene-selective ROI, the parahippocampal place area or PPA. In the first column of Figure 7, the correlation profiles look similar to those in the LOC. In the pairwise comparison in the second column (*residual Layer 7-residual Layer 1*, cyan), we can also observe an initial negative window and a later positive window (although small), indicating an early higher correlation with the *residual Layer 1* and a later higher correlation with *residual Layer 7*. In contrast, in the pairwise comparison in the third column (*residual Layer 1-common components*, yellow), we did not observe any significant differences. In the pairwise comparison in the fourth column (*residual Layer 7-common components*, magenta), we again observe an early negative window and a later positive window, indicating early higher correlation with the object-category-relevant low-level features as compared to the high-level features, and a reversed pattern at later time points. Interestingly, this detected positive window is near 500 ms, perhaps indicating sustained processing of high-level features in PPA.

The fourth row of Figure 7 shows the results in a scene-selective ROI near the transverse occipital sulcus (TOS). In the first column, the correlation profiles appear to be in the middle of a transition from the pattern in the EVC to the patterns of the LOC and the PPA. This may be because the TOS is closer to the EVC than the LOC in euclidean distance in the source space. The early peaks of the blue and red curves look similar to those in the LOC; yet the later peaks starting near 300 ms look similar to those in the EVC, but with much smaller magnitudes. In the pairwise comparison in the second column of Figure 7 (*residual Layer 7-residual Layer 1*, cyan), again, we see an initial negative window and later positive windows, indicating early higher correlation with the *residual Layer 1* and later higher correlation with *residual Layer 7*. The latest significant positive window is at 500 ms, which may indicate some sustained processing of high-level features. In the pairwise comparison in the third column (*residual Layer 1-common components*, yellow), we observe a positive, but not significant, window from about 60 ms to 120 ms, and several negative windows at 170 ms, 260 ms and 320 ms, which indicates a shift of the correlation effect from the less-object-category-relevant low-level features to the object-category-relevant low-level features, consistent with feedforward information flow. In the pairwise comparison in the fourth column (*residual Layer 7-common components*, magenta), we observe an early negative window and a later positive window, indicating early higher correlation with the object-category-relevant low-level features as compared to the high-level features; we also observe a pattern that reverses in later time windows.

The fifth row of Figure 7 shows the results in a scene-selective ROI near the retrosplenial complex (RSC). In the first column, the correlation profiles appear similar to those in the LOC and the PPA. In the pairwise comparison in the second column of Figure 7 (*residual Layer 7-residual Layer 1*, cyan), we see an initial negative window near 60 ms to 120 ms and a later positive window near 170 ms, but no later positive windows were detected in the statistical tests. This pattern is consistent with an early-to-late shift from the less-object-category-relevant low-level features to high-level features. In the pairwise comparison in the third column (*residual Layer 1-common components*, yellow), we observe a positive window near 60 ms to 120 ms, and a negative window at 150 ms. This pattern is consistent with an early-to-late shift from the less-object-category-relevant low-level features to object-category-relevant low-level features. In the pairwise comparison in the fourth column (*residual Layer 7-common components*, magenta), we observe a negative window near 60 ms to 100 ms, which indicates higher correlation effect with object-category-relevant low-level features than with the high-level features in this early time window. In general, the results of these pairwise comparisons are consistent with feedforward information flow.

To summarize these results, using a regression analyses in source space, we obtained spatio-temporal correlation profiles between neural activities and the three groups of features (the *common components*, the *residual Layer 1* and the *residual Layer 7*). By analyzing these profiles, we observed progressive shifts from early (60 ms to 120 ms) to later (after 150 ms) time windows, from lower-level to higher-level features, and from low-level regions to higher-level regions along the hierarchy. These results strongly support a model of visual cortex in which feedforward information flow is intrinsic to perceptual processing. Perhaps more novel is our observation that at a later time window from about 300 ms to 350 ms the correlation of the EVC with the object-category-relevant low-level features (the *common components*) was larger than the correlation with the low-level features that were roughly orthogonal to high-level features (the *residual Layer 1*). This result suggests that a non-feedforward process (e.g., top-down influences) may facilitate EVC in distinguishing between low-level features carrying different kinds of information and representing those low-level features that are object-category-relevant.

## 4 Discussion

One of the advantages of MEG (as compared to EEG or fMRI) is that it allows for measurement of joint spatio-temporal patterns of neural activity. Thus, we are somewhat surprised that the majority of previous work using MEG to correlate neural responses with computer vision features focused solely on temporal patterns (Clarke et al., 2014; Cichy et al., 2016c,a). In line with the standard hierarchical model of primate visual cortex, these prior studies observed an early-to-late shift from lower-level to higher-level features— a result consistent with the results we present here. Additionally, when joint spatio-temporal patterns were considered, Clarke et al. (2014) observed correlations between neural activity and visual/semantic features in source space indicating feedforward information flow—again, consistent with our present results (although Clarke et al. (2014) do not present the full time course for individual ROIs). Using a somewhat different approach, Cichy et al. (2016d) “fused” fMRI and MEG recordings by comparing the representational similarity of visual objects in the two imaging modalities with no reference to externally-derived features; they too observed an early-to-late shift in a feedforward direction of the cortical hierarchy.

What our study adds to this literature are more comprehensive spatio-temporal profiles of visual cortical processing, in particular, because we rely on features derived from a more sophisticated computer vision model (AlexNet) as compared to the model used in (Clarke et al., 2014) (for a comparison of such models, see Yamins et al. (2014)). Moreover, we included a relatively large number of complex, naturalistic scene images (rather than single objects on blank backgrounds as used in Clarke et al. (2014) and Cichy et al. (2016d)). Finally, and perhaps most uniquely, we developed a novel decomposition of low-level and high-level features, and from them derived three orthogonal sets of features: (1) the common components between the low-level and high-level features, which can be interpreted as object-category-relevant low-level features; (2) the residual low-level features that is roughly orthogonal to high-level features, which can be interpreted as object-category-irrelevant low-level features; and (3) the residual object-category-relevant high-level features, where low-level components are partialed out. By comparing the spatio-temporal neural correlation profiles with these three sets, we provided new evidence for non-feedforward processing within visual cortex.

Th e key observation concerns the temporal correlation profiles of the early visual cortex (EVC) with the two sets of low-level features: (1) the object-category-relevant low-level features and (2) the object-category-irrelevant low-level features. In the early neural response to the stimulus images near 60 to 120 ms, the correlation effects with the low feature sets had similar dynamics, peaking around 80 to 100 ms; the correlation effect with the object-category-irrelevant low-level features was stronger. In a later time window starting near 300 ms, which was likely to be the window of neural responses to the disappearance of the stimulus images at 200 ms, the neural correlation profiles were different between the two sets of features — the object-category-relevant low-level features had stronger and longer correlation with the neural activity in the EVC which lasted till 380 ms. If the visual system only has pure feedforward processing, then EVC is unlikely to differentiate between the two sets of features. A likely explanation is that there is feedback information flow in the visual cortex, potentially from higher-level regions to EVC, that “emphasizes” the processing of the object-category-relevant information, or “guides” the EVC to separate object-category-relevant low-level features from other low-level features. Alternatively, such separation may result from self-organizing behavior (e.g, similar to the self-organizing map, Kohonen (1990)) via lateral connections locally in the EVC, which is essentially similar to a local feedback mechanism. In sum, this novel observation provide unique insights on the information flow of processing naturalistic images of scenes.

### Potential pitfalls and future directions

In order to rigorously investigate the nature of feedforward and feedback information flow within the visual cortex we utilized a high-performing CNN model(AlexNet; (Krizhevsky et al., 2012)) to estimate low-and high-level features of the stimuli images. This CNN model was pre-trained over 1,000,000’s of images for a task of visual object categorization. Critically, this categorization task was defined by *human-generated* labels over these images; as such, it is expected that AlexNet does a reasonable job capturing human visual processing of natural images. Moreover, based on our observation of a linear dependence between low-and high-level features as represented in this and related models, we developed a novel method for separating features into three orthogonal sets: object-category-relevant low-level features, object-category-irrelevant low-level features, object-category-relevant high-level features. In tandem with the high temporal resolution and reasonable spatial resolution afforded by MEG, we were able to observe evidence for both bottom-up (feedforward) and potentially top-down (feedback) mechanisms during naturalistic scene perception. However, as with all studies, there are specific limitations in our choice of experimental methods and analyses—next we discuss these challenges as well as relevant future directions.

### Neural responses to the disappearance of stimuli

In our experimental design, each stimulus image was presented for 200 ms and then disappeared—the screen switching back to a fixation cross (“+”) displayed against a gray background. This disappearance necessarily produced changes in the visual input which will drive neural responses in early visual cortex. Liang et al. (2008) characterized the magnitudes of responses in cat V1 to the disappearance of stimuli, which were, somewhat unexpectedly, comparable to the magnitudes of responses to the appearance of stimuli. In our analyses, although the mean responses to the appearance and disappearance of all stimuli were subtracted from the data during preprocessing, the image-specific responses (reflected in deviations from the mean) may be correlated with features for each image. This correlation is one possible explanation for why we observed a late peak near 300 ms in the effects associated with the *common components* and the *residual Layer 1* in the EVC. However, from the perspective of the complete visual processing stream the disappearance of stimuli should not be treated as equivalent to the appearance of stimuli, because in contrast to appearance, with disappearance there is no additional semantic information presented across the change in visual input. Consistent with this logic, the *residual Layer 7*) did not show any increase in correlation effects from 300 ms to 400 ms—the time window where responses to stimulus disappearance are likely to occur.

An alternative view of what leads to the peak near 300 ms associated with low-level features is that, post-stimulus, there is some “replay” of the stimulus. In particular, EVC exhibited a stronger and longer correlation effect with the *common components* (the object-category-relevant low-level features) as compared to the *residual Layer 1* (the low-level features that were roughly orthogonal to high-level features). This “discrimination” between object-category-relevant and object-category-irrelevant features hints that the EVC is—through top-down information flow— guided to “replay” category-relevant low-level features. In contrast, if we assumed only feedforward processing, we would expect similar correlation effects for both groups of low-level features (as seen in the early correlation effects near 100 ms in the first EVC plot of Figure 7)— however, in the later time window we instead observe a distinction in EVC between the two feature sets. We posit that a non-feedforward processes, for example, feedback from the higher-level cortex to the EVC or lateral recurrent interactions within the EVC neurons, provide a better explanation of our pattern of results.

Note that feedback-driven visual “replay” may evoke the experience of a visual aftereffect that is perceivable to participants. Hence our results provide some insight for further tests of the above speculation via behavioral experiments. We hypothesize that if participants perceive a stimulus aftereffect due to top-down information flow (as opposed to, for example, a retinal aftereffect), then this aftereffect should be mainly driven by the object-category-relevant low-level features. As such, one could design stimuli to manipulate presence/absence of such features, thereby manipulating the degree of the aftereffect.

As a related point, our design used a relatively short presentation duration (200 ms) in order to reduce saccade artifacts; additionally, we did not mask our stimuli post presentation (e.g., using white noise patterns). The above challenges may be considered as a result of these design limitations, where the disappearance of the stimulus images interfered with the intact dynamics of visual processing. However, the positive here is that our study also provides novel observations regarding feedback within the visual system—the specifics of which have not been discussed much in the literature as far as we are aware. Consequently, future experiments should examine how the differential coding of different types of low-level features (*common components* and *residual Layer 1*) change under varying stimulus durations and masking conditions.

### Local contrast

In the canonical correlation analysis of the intact features from `AlexNet`, we observed that CCA components between Layers 1 and 7 have high correlations with local contrast features (not documented in the main text). Interestingly, in our first pass analysis of the data, we did not partial out the local contrast features when obtaining the three groups (the *residual Layer 1*, the *common components* and the *residual Layer 7*). Under these conditions, we obtained neural correlation effects for the *common components* that were much higher than those for either the *residual Layer 1* or the *residual Layer 7*, presumably due to a large correlation with local contrast features. In this sense, local contrast appears to be a confounding factor. Nevertheless, local contrast may be inherently included in the statistical regularities of natural images. Intuitively, local receptive fields with high contrast often contain informative features with respect to shape and boundaries— consequently they are typically related to semantic information. Further tests of this hypothesis, which can be implemented with a large set of images, may help us better understand the image statistics of natural images. Moreover, in future studies, to alleviate the influence of this confounding factor, we can design new stimuli by adding high contrast features that are less relevant to semantic information, for example, irregular shadow contours. It would be interesting to study whether such high-contrast features are coded differently from the genuinely informative high-contrast features (e.g., true physical edges or contours) within visual cortex, as well as in a CNN trained on “standard” natural images. Moreover, one might also use a generative neural network (Goodfellow et al., 2014) to create experimental test stimuli that are similar in local contrast and other low-level features to natural images but that do not contain identifiable objects.

### Confounding factors in data-driven experiments

As a coda to our discussion of local contrast as a confounding factor we note that there may exist other image properties that show statistical regularity across natural images. As such, these properties may exhibit significant correlations with neural responses, but be distributed unevenly across the three groups of features we identified in our study (i.e., the *common components, residual Layer 1, residual Layer 7* features). This is a limitation of using natural visual stimuli, where the distribution of features is not an easy manipulable variable. At the same time, in that the visual world typically gives rise to a large dimensional and complicated feature space, it is difficult to form good hypotheses that capture this entire space from scratch. In this context, we view the data-driven exploration of visual processing presented here as an initial step for forming new predictions for future hypothesis-driven experiments.

### Choice of using AlexNet

We used an 8-layer convolutional neural network (CNN; `AlexNet`), which was trained to classify images into 1,000 object categories. In contrast, although our stimuli—natural images of scenes—contained objects that fell into these 1,000 categories, it is unlikely that our participants processed them only in this limited way. That is, participants were likely to automatically invoke scene recognition and scene understanding mechanisms during the experiment, although we did not explicitly instruct them to do so. In preliminary data analyses (not presented here), we used features derived from a network of the same architecture as `AlexNet`, but trained to classify 250 scene categories (Zhou et al., 2014b), many of which overlapped with the categories in our stimuli. In this instance, the neural correlation effects we observed were not significantly higher than those with features from `AlexNet`, which was trained on object, rather than scene, categorization. This finding is consistent with a growing body of results suggesting that the features arising in CNNs trained to perform object categorization (as in `AlexNet`) have good transferability to other visual tasks, as suggested by Yosinski et al. (2014) and Huh et al. (2016). One possible explanation is that many of the learned features well characterize broader—task independent—statistical regularities of the visual world. At the same time, scenes contain many object components, and thus scene understanding may benefit from robust mid-level and high-level representations of objects. Supporting this idea, Zhou et al. (2014a) demonstrated that object detectors emerged in a CNN that was trained on scene classification. Conversely, object categorization requires “partialling out” variability arising from the different scenes in which a given object may appear— to the extent that particular scenes are consistent across different objects, these scenes may be learned as a route to more effective object invariance. Therefore, it is not entirely surprising that object-category relevant features as instantiated in a CNN can account for neural data during scene processing.

The rapid pace of progress in AI also places limitations on even recently run studies using what become quite quickly, less-than-state-of-the-art models. Since the publication of `AlexNet`, deeper, more sophisticated, and higher performing CNNs have been developed for a more diverse set of vision tasks. Although it will be interesting to use features from these new networks to explain neural data, we do not believe that using one of many newer models would qualitatively change our results. First, as discussed above, the features learned by `AlexNet` are already very rich and expressive. Second, practically speaking, the number of images we can present in a neuroimaging study will typically be fewer than ~10^3^; limited by such numbers, it is difficult to imaging a large advantage for more sophisticated networks. Given the evidence we found of non-feedforward processing in visual cortex, we hold that it would be more profitable, in future work, to develop and use non-feedforward neural networks, relying on feedback isomorphic to the connectivity structure in the brain (Yu et al., 2016). In this way, we can compare the dynamics of a high-performing network with spatio-temporal neural activity in the brain so as to better understand the information flow.

### Limited number of observations and choosing linear regression

As note, neuroimaging methodologies impose inherent limits on the number of stimuli that may be used: here we presented only 362 scene images. Compared with the high-dimensional features instantiated in `AlexNet`, we did not have a sufficient number of observations to fit large regression models that included all dimensions in the features, or to use higher-capacity and non-linear models. In future work, it is important to vastly increase the number of observations (as in (Chang et al., 2018)), by increasing data collection time or, perhaps, by reducing the number of image repetitions but increasing measurement SNR.

We used linear regression and lower dimensionality for the derived feature sets primarily due to the limitation of data size. One alternative is to rely on representational similarity analysis (RSA) instead of linear regression to test non-linear dependence. However, for RSA and similar types of non-parametric independence tests (e.g. (Gretton et al., 2005)), the testing statistics are mainly useful with permutations to test if significant non-linear dependence is present. The statistics themselves are quite sensitive to how many dimensions are used and how redundant the dimensions are in the neural responses and the features. For example, in the feature set, if one dimension that is highly correlated with the neural responses is duplicated, we may obtain higher statistics in RSA. In contrast, a regression analysis will yield similar results if the different dimensions in the regressors are orthogonalized. In this sense, using the currently existing RSA-like analysis may not be the best way to compare neural dependence with different sets of features.

### Limitations of the spatial resolution of MEG

Finally, it is worth pointing out that although our study focuses on joint spatio-temporal patterns of neural activity, the spatial resolution of MEG is inherently limited by the underdetermined nature of the source localization problem. That is, in MEG there exist a limited number of sensors, but there are many more source points in the brain. Moreover, the spatial correlations in the columns of the forward matrix can produce a spatial blurring effect in the reconstructed source solutions. Both these factors add uncertainty in the localization. Although we applied a widely used dSPM method—as well as a sparsity inducing source-space regression method (Yang et al., 014) as discussed in the Supplementary Materials—there is likely a fundamental limit that affects both localization methods. In particular, because of the “underdeterminedness”, we had to exploit some priors, but the true neural activity may violate these assumptions. Consequently, although we have tried our best to make reasonable assumptions while keeping the models tractable, there is some possibility that our source-space results may deviate from the underlying truth. We hope the future development of theories of source localization and the sensor density of MEG (or EEG, see (Robinson et al., 2017)), as well as experimental work with intracranial recordings in human patients and animals can further validate our findings.

## 5 Conclusions

In this paper, we present detailed spatio-temporal correlation profiles of neural activity using different feature sets derived from the hierarchically-organized layers of an 8-layer CNN—a high-performing vision model for object categorization. Uniquely, we derived three sets of features from the Layer 1 and Layer 7 features of the CNN, which represented the object-category-relevant low-level features (the *common components*), the low-level features roughly orthogonal to high-level features (the *residual Layer 1*), and the unique high-level features that were roughly orthogonal to low-level features (the *residual Layer 7*). Our correlation profiles across these feature sets indicated an early-to-late shift from lower-level features to higher-level features and from low-level regions to higher-level regions along the cortical hierarchy, consistent with standard models of feedforward information flow. Moreover, by contrasting the correlation effects of the *common components* and the *residual Layer 1*, we found that early visual cortex showed a higher and longer correlation effect with the *common components* (the object-category-relevant low-level features) than with the *residual Layer 1* (the low-level features roughly orthogonal to high-level features), in a later time window, possibly in responses to the disappearance of the stimuli. This temporally late, but spatially early, distinction between the two types of low-level features suggests that some non-feedforward processes, such as top-down influences, appears to facilitate the early visual cortex in differentially representing object-category-relevant and object-category-irrelevant information.

## Acknowledgements

This work was supported by 1439237 NSF and RO1 MH64537/MH/NIMH NIH HHS/United States. The author Ying Yang was supported by the Henry L. Hillman Presidential Fellowship at Carnegie Mellon University. We thank Kevin Tan and Austin Marcus for their help in data collection.

## References

Aminoff, E. M., Toneva, M., Shrivastava, A., Chen, X., Misra, I., Gupta, A., and Tarr, M. J. (2015). Applying artificial vision models to human scene understanding. Frontiers in computational neuroscience, 9.

Bar, M. and Aminoff, E. (2003). Cortical analysis of visual context. Neuron, 38(2):347–358.

Bar, M., Kassam, K. S., Ghuman, A. S., Boshyan, J., Schmid, A. M., Dale, A. M., Hämäläinen, M. S., Marinkovic, K., Schacter, D. L., Rosen, B. R., et al. (2006). Top-down facilitation of visual recognition. Proceedings of the National Academy of Sciences of the United States of America, 103(2):449–454.

Benjamini, Y. and Hochberg, Y. (1995). Controlling the false discovery rate: A practical and powerful approach to multiple testing. Journal of the Royal Statistical Society, 57:289–300.

Chang, N. C., Aminoff, E. M., Pyles, J. A., Tarr, M. J., and Gupta, A. (2018). Scaling Up Neural Datasets: A public fMRI dataset of 5000 scenes. In Vision Sciences Society, St. Pete Beach, FL.

Chen, X., Shrivastava, A., and Gupta, A. (2013). Neil: Extracting visual knowledge from web data. In Proceedings of the IEEE International Conference on Computer Vision, pages 1409–1416.

Cichy, R. M., Khosla, A., Pantazis, D., and Oliva, A. (2016a). Dynamics of scene representations in the human brain revealed by magnetoencephalography and deep neural networks. NeuroImage.

Cichy, R. M., Khosla, A., Pantazis, D., Torralba, A., and Oliva, A. (2016b). Comparison of deep neural networks to spatio-temporal cortical dynamics of human visual object recognition reveals hierarchical correspondence. Scientific reports, 6.

Cichy, R. M., Khosla, A., Pantazis, D., Torralba, A., and Oliva, A. (2016c). Deep neural networks predict hierarchical spatio-temporal cortical dynamics of human visual object recognition. arXiv preprint arXiv:1601.02970.

Cichy, R. M., Pantazis, D., and Oliva, A. (2016d). Similarity-based fusion of meg and fmri reveals spatio-temporal dynamics in human cortex during visual object recognition. Cerebral Cortex, page bhw135.

Clarke, A., Devereux, B. J., Randall, B., and Tyler, L. K. (2014). Predicting the time course of individual objects with meg. Cerebral Cortex, page bhu203.

Cox, R. W. (1996). Afni: software for analysis and visualization of functional magnetic resonance neuroimages. Computers and Biomedical research, 29(3):162–173.

Dale, A. M., Liu, A. K., Fischl, B. R., Buckner, R. L., Belliveau, J. W., Lewine, J. D., and Halgren, E. (2000). Dynamic statistical parametric mapping: combining fmri and meg for high-resolution imaging of cortical activity. Neuron, 26(1):55–67.

Deng, J., Dong, W., Socher, R., Li, L.-J., Li, K., and Fei-Fei, L. (2009). Imagenet: A large-scale hierarchical image database. In Computer Vision and Pattern Recognition, 2009. CVPR 2009. IEEE Conference on, pages 248–255. IEEE.

DiCarlo, J. J. and Cox, D. D. (2007). Untangling invariant object recognition. Trends in cognitive sciences, 11(8):333–341.

Engel, S. A., Glover, G. H., and Wandell, B. A. (1997). Retinotopic organization in human visual cortex and the spatial precision of functional mri. Cerebral cortex, 7(2):181–192.

Epstein, R., Harris, A., Stanley, D., and Kanwisher, N. (1999). The parahippocampal place area: Recognition, navigation, or encoding? Neuron, 23(1):115–125.

Fellbaum, C. (1998). Wordnet: An electronic lexical database: Bradford book.

Felleman, D. J. and Van Essen, D. C. (1991). Distributed hierarchical processing in the primate cerebral cortex. Cerebral cortex, 1(1):1–47.

Fischl, B., Salat, D. H., Busa, E., Albert, M., Dieterich, M., Haselgrove, C., Van Der Kouwe, A., Killiany, R., Kennedy, D., Klaveness, S., et al. (2002). Whole brain segmentation: automated labeling of neuroanatomical structures in the human brain. Neuron, 33(3):341–355.

Freeman, J., Ziemba, C. M., Heeger, D. J., Simoncelli, E. P., and Movshon, J. A. (2013). A functional and perceptual signature of the second visual area in primates. Nature neuroscience, 16(7):974–981.

Goodfellow, I., Pouget-Abadie, J., Mirza, M., Xu, B., Warde-Farley, D., Ozair, S., Courville, A., and Bengio, Y. (2014). Generative adversarial nets. In Advances in Neural Information Processing Systems, pages 2672–2680.

Gramfort, A., Luessi, M., Larson, E., Engemann, D. A., Strohmeier, D., Brodbeck, C., Parkkonen, L., and Hämäläinen, M. S. (2014a). MNE software for processing MEG and EEG data. Neuroimage, 86:446–460.

Gramfort, A., Luessi, M., Larson, E., Engemann, D. A., Strohmeier, D., Brodbeck, C., Parkkonen, L., and Hmlinen, M. S. (2014b). Mne software for processing meg and eeg data. NeuroImage, 86(0):446 – 460.

Gretton, A., Bousquet, O., Smola, A., and Schölkopf, B. (2005). Measuring statistical dependence with hilbert-schmidt norms. In International conference on algorithmic learning theory, pages 63–77. Springer.

Grill-Spector, K., Kushnir, T., Edelman, S., Avidan, G., Itzchak, Y., and Malach, R. (1999). Differential processing of objects under various viewing conditions in the human lateral occipital complex. Neuron, 24(1):187–203.

Hamalainen, M., Hari, R., Ilmoniemi, R. J., Knuutila, J., and Lounasmaa, O. V. (1993). Magnetoencephalography–theory, instrumentation, to noninvasive studies of the working human brain. Reviews of Modern Physics, 65:414–487.

Hubel, D. H. and Wiesel, T. N. (1968). Receptive fields and functional architecture of monkey striate cortex. The Journal of physiology, 195(1):215–243.

Huh, M., Agrawal, P., and Efros, A. A. (2016). What makes imagenet good for transfer learning? arXiv preprint arXiv:1608.08614.

Ito, M. and Komatsu, H. (2004). Representation of angles embedded within contour stimuli in area v2 of macaque monkeys. The Journal of neuroscience, 24(13):3313–3324.

Jia, Y., Shelhamer, E., Donahue, J., Karayev, S., Long, J., Girshick, R., Guadarrama, S., and Darrell, T. (2014). Caffe: Convolutional architecture for fast feature embedding. In Proceedings of the 22nd ACM international conference on Multimedia, pages 675–678. ACM.

Khaligh-Razavi, S.-M. and Kriegeskorte, N. (2014). Deep supervised, but not unsupervised, models may explain it cortical representation. PLoS Comput Biol, 10(11):e1003915.

Kohonen, T. (1990). The self-organizing map. Proceedings of the IEEE, 78(9):1464–1480.

Krizhevsky, A., Sutskever, I., and Hinton, G. E. (2012). Imagenet classification with deep convolutional neural networks. In Advances in neural information processing systems, pages 1097–1105.

LeCun, Y., Bengio, Y., and Hinton, G. (2015). Deep learning. Nature, 521(7553):436–444.

Leeds, D. D., Seibert, D. A., Pyles, J. A., and Tarr, M. J. (2013). Comparing visual representations across human fMRI and computational vision. J Vis, 13(13):25.

Liang, Z., Shen, W., Sun, C., and Shou, T. (2008). Comparative study on the offset responses of simple cells and complex cells in the primary visual cortex of the cat. Neuroscience, 156(2):365–373.

Maris, E. and Oostenveld, R. (2007). Nonparametric statistical testing of EEG-and MEG-data. Journal of neuroscience methods, 164(1):177–190.

Nestor, A., Vettel, J. M., and Tarr, M. J. (2008). Task-specific codes for face recognition: how they shape the neural representation of features for detection and individuation. PloS one, 3(12):e3978.

Robinson, A. K., Venkatesh, P., Boring, M. J., Tarr, M. J., Grover, P., and Behrmann, M. (2017). Very high density EEG elucidates spatiotemporal aspects of early visual processing. Scientific Reports, 7(1):16248.

Russakovsky, O., Deng, J., Su, H., Krause, J., Satheesh, S., Ma, S., Huang, Z., Karpathy, A., Khosla, A., Bernstein, M., et al. (2015). Imagenet large scale visual recognition challenge. International Journal of Computer Vision, 115(3):211–252.

Ségonne, F., Dale, A., Busa, E., Glessner, M., Salat, D., Hahn, H., and Fischl, B. (2004). A hybrid approach to the skull stripping problem in mri. Neuroimage, 22(3):1060–1075.

Tanaka, K. (1996). Inferotemporal cortex and object vision. Annual review of neuroscience, 19(1):109–139.

Wasserman, L. (2010). All of Statistics: A Concise Course in Statistical Inference. Springer Publishing Company, Incorporated.

Xu, Y., Sudre, G. P., Wang, W., Weber, D. J., and Kass, R. E. (2011). Characterizing global statistical significance of spatiotemporal hot spots in magnetoencephalography/electroencephalography source space via excursion algorithms. Statistics in medicine, 30(23):2854–2866.

Yamins, D. L., Hong, H., Cadieu, C. F., Solomon, E. A., Seibert, D., and DiCarlo, J. J. (2014). Performance-optimized hierarchical models predict neural responses in higher visual cortex. Proceedings of the National Academy of Sciences, 111(23):8619–8624.

Yamins, D. L. K. and DiCarlo, J. J. (2016). Using goal-driven deep learning models to understand sensory cortex. Nat Neurosci, 19(3):356–365.

Yang, Y. (2017). Source-Space Analyses in MEG/EEG and Applications to Explore Spatio-temporal Neural Dynamics in Human Vision. PhD dissertation, Carnegie Mellon University.

Yang, Y., Tarr, M. J., and Kass, R. E. (2014). Estimating learning effects: A short-time fourier transform regression model for MEG source localization”. In Lecture Notes on Artificial Intelligence: MLINI 2014: Machine learning and interpretation in neuroimaging, Montreal, Canada. Springer.

Yosinski, J., Clune, J., Bengio, Y., and Lipson, H. (2014). How transferable are features in deep neural networks? In Advances in neural information processing systems, pages 3320–3328.

Yu, C.-P., Maxfield, J., and Zelinsky, G. J. (2016). Generating the Features for Category Representation using a Deep Convolutional Neural Network. In Vision Sciences Society, page 1161876.

Zeiler, M. D. and Fergus, R. (2014). Visualizing and understanding convolutional networks. In European conference on computer vision, pages 818–833. Springer.

Zhou, B., Khosla, A., Lapedriza, A., Oliva, A., and Torralba, A. (2014a). Object detectors emerge in deep scene cnns. arXiv preprint arXiv:1412.6856.

Zhou, B., Lapedriza, A., Xiao, J., Torralba, A., and Oliva, A. (2014b). Learning deep features for scene recognition using places database. In Advances in neural information processing systems, pages 487–495.

